# Focus on the spectra that matter by clustering of quantification data in shotgun proteomics

**DOI:** 10.1101/488015

**Authors:** Matthew The, Lukas Käll

## Abstract

In shotgun proteomics, the information extractable from label-free quantification experiments is typically limited by the identification rate and the noise level in the quantitative data. This generally causes a low sensitivity in differential expression analysis on protein level. Here, we propose a quantification-first approach for peptides that reverses the classical identification-first workflow. This prevents valuable information from being discarded prematurely in the identification stage and allows us to spend more effort on the identification process. Specifically, we introduce a method, Quandenser, that applies unsupervised clustering on both MS1 and MS2 level to summarize all analytes of interest without assigning identities. Not only does this eliminate the need for redoing the quantification for each new set of search parameters and engines, but it also reduces search time due to the data reduction by MS2 clustering. For a dataset of partially known composition, we could now employ open modification and *de novo* searches to identify analytes of interest that would have gone unnoticed in traditional pipelines. Moreover, Quandenser reports error rates for feature matching, which we integrated into our probabilistic protein quantification method, Triqler. This propagates error probabilities from feature to protein level and appropriately deals with the noise in quantitative signals caused by false positives and missing values. Quandenser+Triqler outperformed the state-of-the-art method MaxQuant+Perseus, consistently reporting more differentially abundant proteins at 5% FDR: 123 vs. 117 true positives with 2 vs. 25 false positives in a dataset of partially known composition; 62 vs. 3 proteins in a bladder cancer set; 8 vs. 0 proteins in a hepatic fibrosis set; and 872 vs. 661 proteins in a nanoscale type 1 diabetes set. Compellingly, in all three clinical datasets investigated, the differentially abundant proteins showed enrichment for functional annotation terms.

The source code and binary packages for all major operating systems are available from https://github.com/statisticalbiotechnology/quandenser, under Apache 2.0 license.

## Introduction

In mass spectrometry-based proteomics, label-free quantification (LFQ) is one of the most comprehensive methods to analyze protein concentrations in complex mixtures. Its main advantage is that it allows for comparisons in large sample cohorts and can, hence, handle complex experimental designs [2]. Currently, LFQ and quantitative proteomics in general, are struggling to obtain sufficient coverage of the proteome [3] and also suffer from low sensitivity for differentially abundant proteins at false discovery rate thresholds [33]. While this can partially be attributed to inherent limitations in the methodology of mass spectrometry, it is, to a high degree, caused by the inadequacy of our current data analysis pipelines. We also note that LFQ is sometimes seen as cumbersome, as contrary to, for instance, isobaric labeling, one is not guaranteed a readout for an identified peptide in each sample. Frequently, this is resolved by missing value imputation but this introduces a multitude of issues [45, 22]. Novel methods for LFQ data analysis are necessary to address these problems in sensitivity and specificity.

Two well-recognized issues regarding the sensitivity of LFQ pipelines are that many MS1 features remain unassigned to peptides [46] or even to MS2 spectra [29] and that a large number of fragment spectra remain unidentified [37]. Match-between-runs (MBR) has proven to be an effective technique to propagate MS2 information to the MS1 features [5, 46, 1] and clustering of MS2 spectra from large repositories has allowed us to zoom in on frequently unidentified spectra for peptide identification [13]. Unsupervised clustering of MS2 spectra also significantly reduces the number of MS2 spectra that need to be searched [10, 12, 39]. This allows more computationally expensive searches to be conducted such as partial digestions, variable modification searches, open modification searches or even *de novo* searches. Generally, clustering of fragment spectra was recently demonstrated to give better sensitivity to LFQ experiments [14].

A less highlighted issue concerning sensitivity is that conventional data analysis pipelines match the individual sample’s spectra to peptide sequences with search engines as a first step [5, 46]. This step directly limits the number of peptides and proteins that can ultimately be quantified. Moreover, such an *identification-first* strategy has some undesirable properties. For instance, if one would want to search the data again with an open modification search engine or a newly acquired spectral library, all quantification would have to be redone. Yet, in reality, the underlying experimental quantitative data does not change due to a new search engine result, just our interpretation thereof. By using unsupervised clustering of MS1 features, we can capture this underlying quantitative information of the analytes. We can assign MS2 spectra to such clusters but postpone their interpretation to a later stage. Such a *quantification-first* approach for peptides allows us to focus our efforts on improving the coverage by investigating frequently unidentified features without having to go through the costly process of repeated iterations of identification and quantification [44, 23, 31, 9].

We note that the term quantification-first refers to the quantification of peptides, as the quantification of proteins requires some form of identification. Furthermore, current quantification-first approaches often apply significance filters for the differentially abundant peptide signals before peptide and protein identification [31, 9]. However, we refrain from applying such a filter, as this can easily introduce quantification biases on protein level because it discards evidence in support of the null hypothesis. Whenever practically possible, significance tests should be withheld until the final steps of any data processing pipeline, as they introduce unnecessary bifurcations, resulting in loss of data.

Conversely, specificity is often compromised by the multitude of thresholds in the various steps of a quantification pipeline, which cause a lack of error control and do not properly account for error propagation [40]. Also, missing value imputation is known to induce false quantification accuracy [16]. We have previously introduced a hirarchical Bayesian model [], Triqler [40], for protein quantification that propagates error rates of the different steps in protein quantification and compensates for missing readouts in a Bayesian manner. Triqler provides a natural means to link the quantification to the identification process. However, two features that Triqler did not include yet were the error rates of the association of MS1 features with MS2 spectra and support for MBR.

Here, we introduce a new method, *Quandenser* (QUANtification by Distillation for ENhanced Signals with Error Regulation), that we subsequently interface to Triqler to substantially increase the sensitivity of the LFQ analysis pipeline. Quandenser condenses the quantification data by applying unsupervised clustering on both MS1 and MS2 level and thereby paves the way for a quantification-first approach. The algorithm combines the unknowns from both levels into a condensed set of MS1 features and MS2 spectra that are likely to be interesting for further investigation. Specifically, Quandenser incorporates a method similar to match-between-runs for MS1 features to increase sensitivity and uses MS2 spectrum clustering to decrease the number of spectra to be searched. Importantly, Quandenser addresses the false transfer problem [24] by providing *feature-feature match* error rates using decoy features and a novel automated weighting scheme to separate true from false matches. These error rates can be used as input to Triqler to account for the errors as a result of the MBR step.

The main advancement of our pipeline of Quandenser with Triqler comes from the combination of a match-between-runs approach with a Bayesian error model, which results in compelling gains in the number of significant proteins while maintaining control over the differential abundance FDR. Additionally, the clustering approach provides a conceptually pleasing solution to some of the bottlenecks in protein quantification analysis by producing a reduced set of hypotheses to test for. Not only does this facilitate easy re-searching and open modification searches on the data, but it also allows us to zoom in on previously unexplored parts of our data that, for example, are frequently recurring but remain consistently unidentified or follow a quantitative behavior of potential interest.

## Methods

### Datasets

We downloaded RAW files for 4 engineered datasets to characterize the reliability and sensitivity of our pipeline. The first dataset was a set of partially known composition with the UPS1 protein mixture spiked in at different concentrations in a yeast background (PRIDE project: PXD002370, 9 RAW files, Only the files C-25fmol-R* QEx2 *, D-10fmol-R* QEx2 *, and E-5fmol-R* QEx2 * were used) [11]. The other 3 sets were mixtures of two proteomes spiked in at different ratios. The first comes from a benchmark comparing quantitative accuracy and coverage between the commonly used LFQ and TMT methods using a sample of yeast spiked into a human background [32] (PRIDE project: PXD007683, 11 LFQ RAW files). Second, we analyzed the data from a benchmark for MS1 label-free quantification where E. coli was spiked into a HeLa background at 4 concentrations [35] (PRIDE project: PXD001385, 12 RAW files), which will be referred to as the *Shalit hela-ecoli* dataset. Finally, we processed the data from the recent BoxCar method, where again E. coli was spiked into a HeLa background at two different ratios [28] (PRIDE project: PXD006109, 12 RAW files, only the files for the mixture of E. coli with HeLa were considered, both from the Shotgun and the BoxCar runs), which will be referred to as the *BoxCar hela-ecoli* dataset.

We also downloaded RAW files for three clinical datasets. The first was a dataset studying bladder cancer [21] (PRIDE project: PXD002170, 8 LFQ RAW files), which will henceforth be referred to as the *Latosinska* dataset. The second was a dataset studying Hepatitis C Virus-associated hepatic fibrosis [4] (PRIDE project: PXD001474, 27 RAW files), which will be referred to as the *Bracht* dataset. The third dataset concerned a recent advancement in nanoscale proteomics applied to type 1 diabetes [48] (PRIDE project: PXD006847, 18 RAW files), which will be referred to as the *Zhu* dataset.

For the UPS-yeast mixture, a UPS1 protein mixture was spiked into a 1 *µ*g yeast background at respectively 25, 10 and 5 fmol concentration, with triplicates for each concentration. For the human-yeast mixture, yeast lysate was spiked in at 10% (*n* = 3), 5% (*n* = 4) and 3.3% (*n* = 4) total protein concentration into a human cell lysate (SH-SY5Y) background. For the Shalit hela-ecoli mixture, 3, 7.5, 10 and 15 ng of E.coli lysate was spiked into a HeLa digest background of 200 ng in triplicates. For the BoxCar hela-ecoli mixture, E. coli lysate was mixed in 1 : 2 and 1 : 12 ratios in a HeLa lysate in triplicates. The Latosinska dataset featured 8 samples of tumor tissues of non-muscle invasive (stage pTa, *n* = 4) and muscle-invasive bladder cancer cases (stage pT2+, *n* = 4), without technical replicates. The Bracht dataset featured 27 samples of biopsies from patients with HCV-associated hepatic fibrosis, classified in a low fibrosis group (*n* = 13) and a high fibrosis group (*n* = 14), without technical replicates. The Zhu dataset featured 18 samples with 10–100 cells each from human pancreatic islet thin sections with nine samples from a type 1 diabetes donor and nine from a non-diabetic control donor, without technical replicates.

Prior to processing the runs with Quandenser v0.02, all RAW files were converted to mzML format with ProteoWizard [18] (ver 3.0.10765), where we applied peak picking both on MS1- and MS2-level, except for the Bracht set, where peak picking was only applied on MS2-level. For the BoxCar runs of the BoxCar dataset, MS1 BoxCar windows were combined into new MS1 spectra using an in-house python script.

### Quandenser

An overview of the Quandenser process is given in Figure 1. First, MS1 features were detected with Dinosaur v1.1.3 [38] and assigned to the MS2 spectra that were obtained inside the retention time and precursor isolation window. We will refer to a combination of an MS1 feature and an MS2 spectrum as a *spectrum-feature match*. Next, clustering of MS2 spectra was applied with MaRaCluster v0.05 [39]. One advantage of applying MS2 clustering first was that we could align retention times between two runs through pairs of spectra that end up in the same cluster. This alignment was done by fitting a spline function to the medians from 100 bins of the sorted retention times using iteratively reweighted least squares regression (IRLS). The IRLS algorithm protected against outliers that might have resulted from incorrect clusterings. We then estimated the standard deviation of the aligned retention times, which gave us a way to dynamically select a reasonable window to search for matching precursors (by default, 5 standard deviations), instead of having this window specified with a fixed time interval, as is usually done.

**Figure 1:**
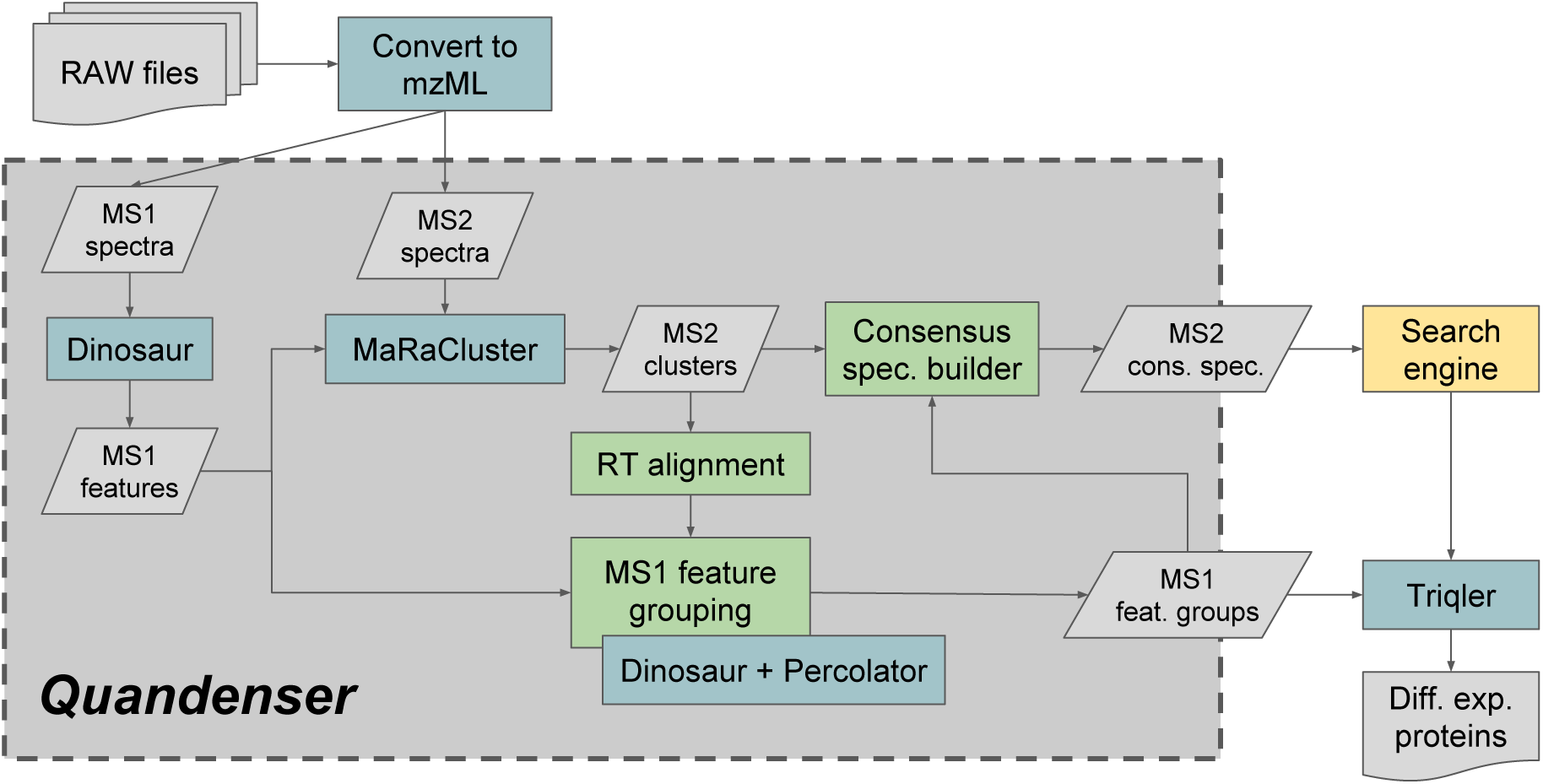
Workflow within Quandenser and its employment in the protein quantification pipeline. Blue boxes represent existing software packages, some of which are directly integrated into the Quandenser package, green boxes represent in-house packages, gray boxes represent intermediate results or files and the yellow box represents any search engine(s) of choice.

Retention times were aligned pair-wise between runs using a minimum spanning tree based on the similarity of chromatography runs [34, 5]. This tree was traversed twice, once from bottom to top and a second time from top to bottom. This ensured that MS1 feature information was distributed between all pairs of runs, not only the pairs connected in the tree. For each pair-wise retention time alignment, the MS1 features discovered by Dinosaur from the two corresponding runs were matched based on a set of numeric features, such as the difference between the observed and aligned retention time and precursor mass difference. At the same time, we matched MS1 features against *decoy MS1 features* [46], which were generated by shifting the precursor m/z of all MS1 features discovered by Dinosaur by 5 · 1.000508 Th in one of the runs. For a more detailed description of these decoy features and what constitutes good decoy features, see Supplementary Section 5. These decoy MS1 features correctly mimicked incorrect target MS1 features in a target-decoy competition setting (Supplementary Figure S10). Also, we noted that the probability of observing a certain precursor mass is constrained by its composition of amino acids and, therefore, the probability of a match quickly tapers off when non-integer multiples of 1.000508 were employed as shifts (Supplementary Figure S10f).

The MS1 feature characteristics, e.g. the precision of precursor m/z and retention time, can vary significantly between samples. We designed Quandenser to allow different amounts of flexibility when matching MS1 features between different runs. We achieved this by automatically weighting the numeric features using Percolator v3.02 [41]. This perhaps surprising use of Percolator - which normally is used for quality assessments of peptide-spectrum matches - allowed us to avoid fixed thresholds for individual numeric features, and instead derive error probabilities for feature-feature matches. This approach is an extension of the approach employed by DeMix-Q [46], where Percolator now provides a more natural means to feature weighting as well as probabilistic scoring.

We retained feature-feature matches with a posterior error probability (PEP) below 0.25, which typically controls the FDR well below 5%. These PEPs were also used as input to Triqler to control for false matching. For features for which no confident corresponding feature could be found in the opposite run, we used a match-between-runs approach by searching for previously missed features using the targeted mode of Dinosaur, again scored by Percolator according to the same scheme as outlined above. If a feature still had no match, a placeholder feature, representing a missing value, was added at the corresponding precursor mass and aligned retention time.

The MS1 features were grouped by feature-feature matches from the pair-wise alignments using single-linkage clustering, resulting in MS1 *feature groups*. The feature groups that had a missing value in more than *M* runs were filtered out. Subsequently, for each feature group, we selected all MS2 spectra that were previously assigned to a feature in the group and assigned the MS2 spectrum’s corresponding cluster from MaRaCluster to the feature group. We then scored each match between a feature group and a spectrum cluster according to the following score:

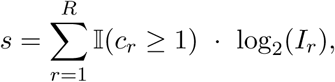

where *r* = 1, …, *R* indicates the run index, 𝕀 is the indicator function that is 1 if the condition holds and 0 otherwise, *c*_*r*_ are the number of spectra in the spectrum cluster that can be linked to the feature for run *r*, and *I*_*r*_ is the intensity of the feature in run *r*. For each spectrum cluster, we only retained the feature groups with 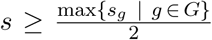, with *G* the set of all feature groups associated with the spectrum cluster. The idea behind this filter is that the MS1 feature intensity is a good predictor of the dominant peptide species in the MS2 spectrum. This filter typically more than halved the number of spectrum-feature matches that had to be searched, while maintaining or even increasing the number of peptide identifications due to the reduction of tested hypotheses.

Finally, Quandenser produces (1) a list of MS1 feature groups with their corresponding MS2 spectrum clusters, and (2) a spectrum file with, for each spectrum cluster, its consensus spectrum together with the spectrum-feature matches of the assigned feature groups. The spectrum file can be any of the file formats that are supported by Proteowizard, such as mzML and mgf, and can then be processed by the user’s search engine of choice.

### Peptide and protein identification and quantification

For the UPS-yeast mixture, we created a concatenated FASTA file containing 6769 records from both the UPS1 proteins (https://www.sigmaaldrich.com/, accessed: 2018 Jan 17) and the Swiss-Prot database for yeast (http://www.uniprot.org/, accessed: 2016 Mar 15). For the human-yeast mixture, we created a concatenated FASTA file with 26914 entries from the Swiss-Prot databases of human (accessed: 2016 Mar 15) and yeast (accessed: 2015 Nov 12). For the Shalit hela-ecoli mixture, we used the FASTA file provided in the PRIDE repository containing 24557 sequences from the Swiss-Prot databases of human (accessed: 2014 Jun) and E. coli (accessed: 2014 Aug). For the BoxCar hela-ecoli mixture, we created a concatenated FASTA file with 24753 proteins of the Swiss-Prot proteins of human (accessed: 2019 Sep 6) and E. coli (accessed: 2019 Apr 11). The Latosinska, Bracht and Zhu sets were searched against the Swiss-Prot database for human containing 20193 records (accessed: 2015 Nov 12). We generated decoy sequences by reversing the amino acid sequences and created concatenated target-decoy databases as input to the search engines.

We used several search engines both as standalone packages as well as part of a cascade search approach [17]. We employed Tide [6], through the interface of the Crux 2.1 [27] package, and used Percolator v3.02 [41] to post-process the resulting PSMs. For all datasets, all parameters in Tide and Percolator were left to their default values (i.e. tryptic cleavages, fixed carbamidomethylations of cysteine and `--mz-bin-width=1.000508`), except for allowing up to 2 methionine oxidations, up to 2 missed cleavages, and ±10 ppm precursor tolerance. Furthermore, we used MODa v1.51 [30] and MSFragger (build 20170103.0) [19] for open modification searches. We extracted several relevant features of the respective search results with in-house python scripts and subsequently processed the PSMs with Percolator v3.02. Finally, we also used Novor v1.05.0573 [26] for *de novo* searches as a discovery tool and searched the resulting sequences with BLAST through the UniProtKB website. We did no statistical analysis on the Novor results.

After the search engine search, the feature groups output file from Quandenser and the search engine results were combined into an input file to Triqler v0.1.4 with a python script available from the Quandenser repository (`bin/prepare_input.py`). This step includes the option for retention time-dependent normalization [46] which is the default option and was also applied to all datasets presented here. The number of missing values allowed for each feature group, *M*, was chosen to reflect the parameters chosen by the authors of the original manuscripts, or, if unavailable, we allowed between 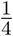 and 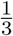 of the runs to have missing values. For the UPS-yeast dataset, we allowed *M* = 3 missing values, for the human-yeast mixture we used *M* = 4, for the Shalit hela-ecoli mixture *M* = 3 and for the BoxCar hela-ecoli set we analyzed all 12 runs together with Quandenser and the respective search engine(s), but then analyzed the 6 shotgun runs and the 6 BoxCar runs separately with Triqler using *M* = 3. As for the clinical datasets, we used *M* = 4 for the Latosinska set, *M* = 7 for the Bracht set and *M* = 11 for the Zhu set. Although the number of allowed missing values has a significant impact on sensitivity, testing different values of *M* on the human-yeast mixture showed that the FDR could still be controlled within reasonable bounds, i.e. below 10% empirical FDR at 5% reported FDR, even when allowing more than 70% of the runs to have missing values (Supplementary Table S32).

The use of Triqler’s hierarchical Bayesian model to combine the quantitative signals of the spectrum-feature matches into protein quantities offers a great way to integrate away undesired variation. The algorithm will in effect down-weight any of a protein’s constituent peptides with a quantification pattern that contradicts its other peptides. In this aspect, Triqler resembles our previously published method, Diffacto [47], which uses factor analysis to obtain a similar effect. However, Triqler expands on this idea with the integration of identification errors, a more intuitive way to impute missing values and posterior probabilities that facilitate better interpretation of the results.

Some adaptations to Triqler were necessary to deal with the additions to the pipeline. Firstly, the feature-feature match PEPs are used explicitly as an extra input to the feature node in Triqler’s probabilistic graphical model (Supplementary Figure S1). Secondly, there is the issue of the many-to-many relation between feature groups and spectrum clusters. Note that a spectrum cluster can be associated with multiple identified peptide sequences due to the chimericity of the spectrum. In the end, only one peptide identification can be associated with each feature group. To resolve this, for each unique peptide sequence identification of the spectrum cluster, Triqler first assigns the feature group with the best search engine score. Subsequently, if a feature group still has multiple peptide identifications, Triqler chooses the peptide sequence with the best combined PEP of the search engine PEP and the feature-feature match PEPs.

Finally, to deal with open modification search results, we have to guard ourselves against assigning correct peptides with incorrect (small) modifications to feature groups. To prevent this, we only select the best peptide identification per protein for each spectrum cluster, under the assumption that the search engine will score the true peptide sequence, with or without modification, the highest. Triqler then proceeds in normal fashion to calculate relative protein expression levels and finally outputs a list of differentially abundant proteins, based on the posterior distributions of the log2 fold change between treatment groups of the protein concentrations.

As a comparison, we also analyzed the datasets with MaxQuant v1.6.1.0 [5], starting from the RAW files, followed by differential expression analysis with Perseus v1.6.1.3 [43]. We used the default parameters in MaxLFQ, except for allowing up to 2 oxidations and allowed the use of these modified peptides for quantification. For the differential expression analysis with Perseus, we filtered out decoy proteins and proteins with more than *M* missing values per dataset as stated above. We then log2 transformed the intensities, used missing value imputation from a down-shifted gaussian with the default parameters and used a two-sided Welch’s t-test with different values of *S*_0_ (0.0, 0.3, 0.7, 1.0), where higher values of *S*_0_ will increasingly prevent small fold changes from being selected as significant [42]. The results reported in the main text are for *S*_0_ = 0.3 unless stated otherwise, as these generally gave the best trade-off between sensitivity and specificity.

Additionally, we analyzed the protein group-level output of MaxQuant with the Empirical Bayesian Random Censoring Threshold model (EBRCT) [20]. This method deals with missing values in a similar way to Triqler, using the assumption that low abundant analytes are more likely to produce missing values. The main difference is that it operates on protein group-level input and is thus a simpler approach. We adapted the R scripts provided in the supplementary materials of the original publication to output the posterior estimations for the group means and the group means’ standard deviations, which we then used to compute fold change posterior distributions.

Finally, we used DAVID 6.8 [15] to find significant functional annotation terms for the sets of differentially abundant proteins found by Quandenser+Triqler and MaxQuant+Perseus. We used the proteins identified at 5% protein-level identification FDR as the background set and thresholded the significant terms at a 5% Benjamin-Hochberg corrected FDR.

## Data availability

RAW files for all 7 datasets analyzed are available without restrictions from the PRIDE repository: UPS-yeast mixture (PXD002370), human-yeast mixture (PXD007683), Shalit hela-ecoli mixture (PXD001385), BoxCar hela-ecoli mixture (PXD006109), Latosinska dataset (PXD002170), Bracht dataset (PXD001474) and the Zhu dataset (PXD006847).

## Code availability

The source code and binary packages for all major operating systems are available from https://github.com/statisticalbiotechnology/quandenser, under Apache 2.0 license. An installation guide and instructions on how to run the full pipeline including Triqler are included in the Supplementary Materials.

## Results

We evaluated the performance of our quantification-first method, Quandenser, with different search engines followed by differential expression analysis with Triqler. We used one regular search engine, Tide, one open modification search engine, MODa and subsequently, we investigated a combination of them both, i.e. Tide and MODa in a cascade search setting. We compared these quantification-first setups to three identification-first setups, (1) MaxQuant+Perseus with match-between-runs (MBR), MaxQuant+EBRCT with MBR and (3) Tide followed by Triqler, without clustering on MS1 and MS2 level nor MBR but with feature detection using Dinosaur. In the following, whenever MaxQuant is mentioned as a method, this is presumed to include MBR.

For the engineered datasets we verified the reported differential abundance FDRs by comparing it to the observed differential abundance FDR. To calculate the latter, spiked-in proteins reported as differentially abundant with the correct fold change sign were counted as true positives. All other proteins reported as differentially abundant, i.e. spiked-in proteins with the wrong fold change sign and background proteins, were counted as false positives.

Furthermore, we used the UPS-yeast dataset to illustrate some benefits of clustering the data before analysis by a search engine. The low complexity of the UPS spike-in fraction allowed us to verify our findings with high confidence. Then, we used the human-yeast proteome mixture to characterize the benefits of each of the steps in our pipeline, as its higher complexity allowed better separation of the performance figures. The two hela-ecoli mixtures were then used to further demonstrate the applicability of the method in different experimental setups. Finally, we analyzed 3 clinical datasets to characterize our pipeline’s behavior in real-world applications. A summary of the datasets, including the employed parameters and reduction in the number of searched MS2 spectra through clustering by Quandenser can be found in Supplementary Table S34.

All analysis was performed on a quad-core (Intel i7-4790K, 4.00GHz) machine with 32 GB of memory running Ubuntu 18.04. Conversion from RAW to mzML format with ProteoWizard took less than 10 minutes per dataset using 4 cores. Both Quandenser and MaxQuant were run using all 4 cores as well. For MaxQuant, we used Mono on Linux, which was reported to be at least as fast as MaxQuant on Windows [36]. On the tested datasets, running Quandenser was typically about twice as fast as MaxQuant, with respective runtimes in the range of 1−3 hours compared to 2−5.5 hours. Quandenser does not include a search engine step, but processing the consensus spectra with Tide and Triqler took less than 15 minutes per dataset. The open modification searches with MODa on the consensus spectra for each of the clinical datasets took about 15 − 18 hours using 4 cores. One should bear in mind that running MODa without applying clustering would have taken between 4 and 6 days.

### UPS-yeast dataset

The UPS-yeast dataset consisted of a total of 535k MS2 spectra. Assigning the MS1 features detected by Dinosaur to the MS2 spectra resulted in 934k spectrum-feature matches. Allowing up to *M* = 3 missing values, resulted in 121k feature groups of which 50k were assigned to at least one spectrum cluster. As multiple spectrum clusters could be assigned to a feature group, 109k spectrum-feature matches (12% of the original number of spectrum-feature matches), corresponding to 61k consensus spectra, remained to be searched by the search engine.

Subsequently, we processed the Quandenser and search engine outputs with Triqler and obtained posterior distributions of the fold changes between sample groups (Supplementary Figure S8). To illustrate what information Triqler uses and how it arrives at these posterior distributions, we have added a detailed example for one of the UPS proteins in Supplementary Section 1.1. Triqler differs from conventional statistical tests in that it integrates the probabilities of the fold change exceeding a given threshold instead of calculating a probability of the observed data given a fold change of zero (a *p* value). For the UPS-yeast dataset, we selected this log2 fold change to be 0.8 (`--fold_change_eval=0.8`), which is just below the lowest spike-in ratio in the 10 vs 5 fmol comparison. Processing the Quandenser output with Triqler resulted in higher sensitivity compared to applying Triqler directly on the search results without clustering and at the same time controlled the FDR below the reported FDR of 5%, whereas MaxQuant+Perseus and MaxQuant+EBRCT failed to do so, producing observed FDRs of up to 26% and 18% respectively (Table 2, Supplementary Table S1). We also observed that MaxQuant+Perseus had trouble controlling the FDR, regardless of the value for the *S*_0_ parameter (Supplementary Table S1).

**Table 1:**
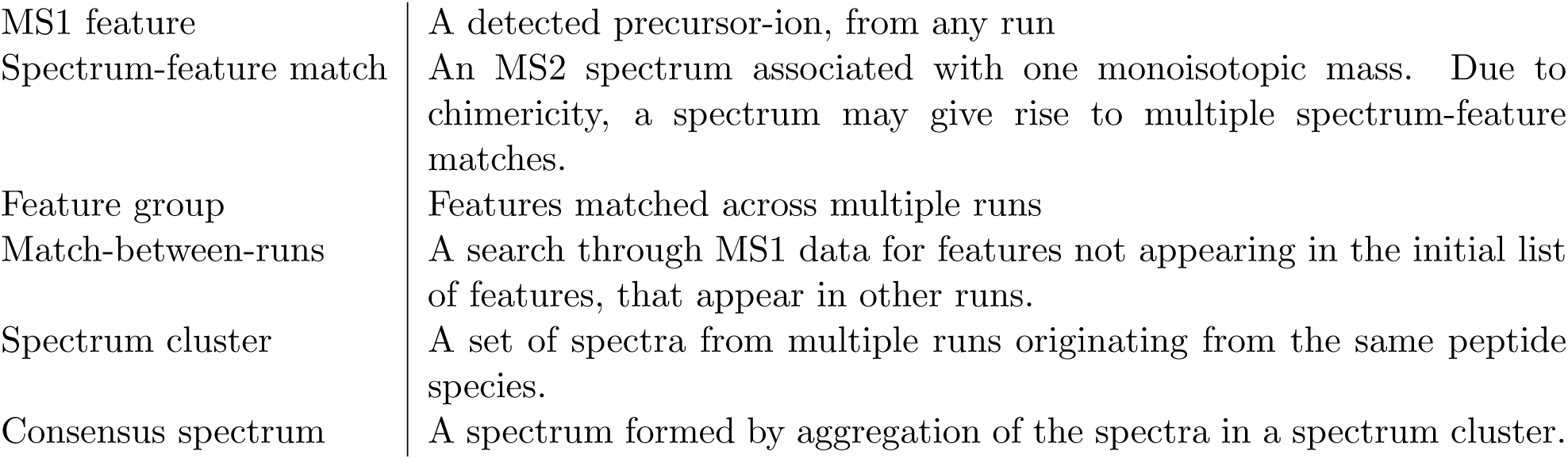
Frequently occurring terms.

|  |  |
| --- | --- |
| MS1 feature | A detected precursor-ion, from any run |
| Spectrum-feature match | An MS2 spectrum associated with one monoisotopic mass. Due to chimericity, a spectrum may give rise to multiple spectrum-feature matches. |
| Feature group | Features matched across multiple runs |
| Match-between-runs | A search through MS1 data for features not appearing in the initial list of features, that appear in other runs. |
| Spectrum cluster | A set of spectra from multiple runs originating from the same peptide species. |
| Consensus spectrum | A spectrum formed by aggregation of the spectra in a spectrum cluster. |

**Table 2:** Quandenser combined with Triqler achieves a high sensitivity on the UPS-yeast and Shalit HeLa+Ecoli datasets, while still maintaining control of the FDR. The table lists the number of true (tp) and false positive (fp) significantly differentially abundant proteins at a 5% reported FDR threshold. For Perseus we used *S*_0_ = 0.3, other values (*S*_0_ = 0.0, 0.7, 1.0) resulted in inferior results (Supplementary Table S1).

UPS-yeast
| UPS1 spike-in concentration [fmol] | 25 vs 10 |  | 25 vs 5 |  | 10 vs 5 |  |
| --- | --- | --- | --- | --- | --- | --- |
| Max. true positives | 48 |  | 48 |  | 48 |  |
| **Method** | tp | fp | tp | fp | tp | fp |
| Quandenser+Triqler |  |  |  |  |  |  |
| - Tide | 43 | 1 | 45 | 1 | 35 | 0 |
| - MODa | 40 | 0 | 41 | 2 | 29 | 0 |
| - Tide + MODa cascade | 43 | 0 | 45 | 0 | 35 | 0 |
| Tide+Triqler (without Quandenser/MBR) | 40 | 0 | 40 | 0 | 34 | 0 |
| MaxQuant MBR+Perseus ($S_0 = 0.3$) | 39 | 9 | 42 | 15 | 36 | 1 |

Shalit hela-ecoli
| E. coli spike-in concentration [ng] | 3 vs 7.5 |  | 3 vs 10 |  | 3 vs 15 |  | 7.5 vs 10 |  | 7.5 vs 15 |  | 10 vs 15 |  |
| --- | --- | --- | --- | --- | --- | --- | --- | --- | --- | --- | --- | --- |
| **Method** | tp | fp | tp | fp | tp | fp | tp | fp | tp | fp | tp | fp |
| Quandenser+Triqler |  |  |  |  |  |  |  |  |  |  |  |  |
| - Tide | 190 | 7 | 198 | 4 | 229 | 0 | 0 | 0 | 194 | 3 | 138 | 0 |
| - MODa | 170 | 3 | 181 | 5 | 222 | 2 | 0 | 0 | 184 | 1 | 131 | 1 |
| - Tide + MODa cascade | 200 | 8 | 206 | 8 | 240 | 3 | 0 | 0 | 201 | 4 | 145 | 1 |
| Tide+Triqler (without Quandenser/MBR) | 47 | 1 | 50 | 0 | 62 | 0 | 0 | 0 | 59 | 1 | 49 | 0 |
| MaxQuant MBR+Perseus ($S_0 = 0.3$) | 126 | 7 | 134 | 6 | 137 | 9 | 0 | 0 | 128 | 4 | 125 | 7 |

To demonstrate the advantages of reducing the number of spectra and spectrum-feature matches that need to be searched, we ran the unidentified consensus spectra through an open modification search with MODa using the cascade search approach [17]. This open modification search took about 4 hours using 4 cores, whereas applying such a search with the same number of cores without clustering by Quandenser would have taken well over a day. We could see a clear increase in the number of feature groups that were assigned a peptide and more modest, but still significant, increases in the number of unique peptides and proteins (Figure 2). However, the cascade search did not result in an increased sensitivity on the spiked-in proteins, as the newly discovered peptides were predominantly modified versions of already identified peptides or came from already identified proteins (Supplementary Table S1).

**Figure 2:**
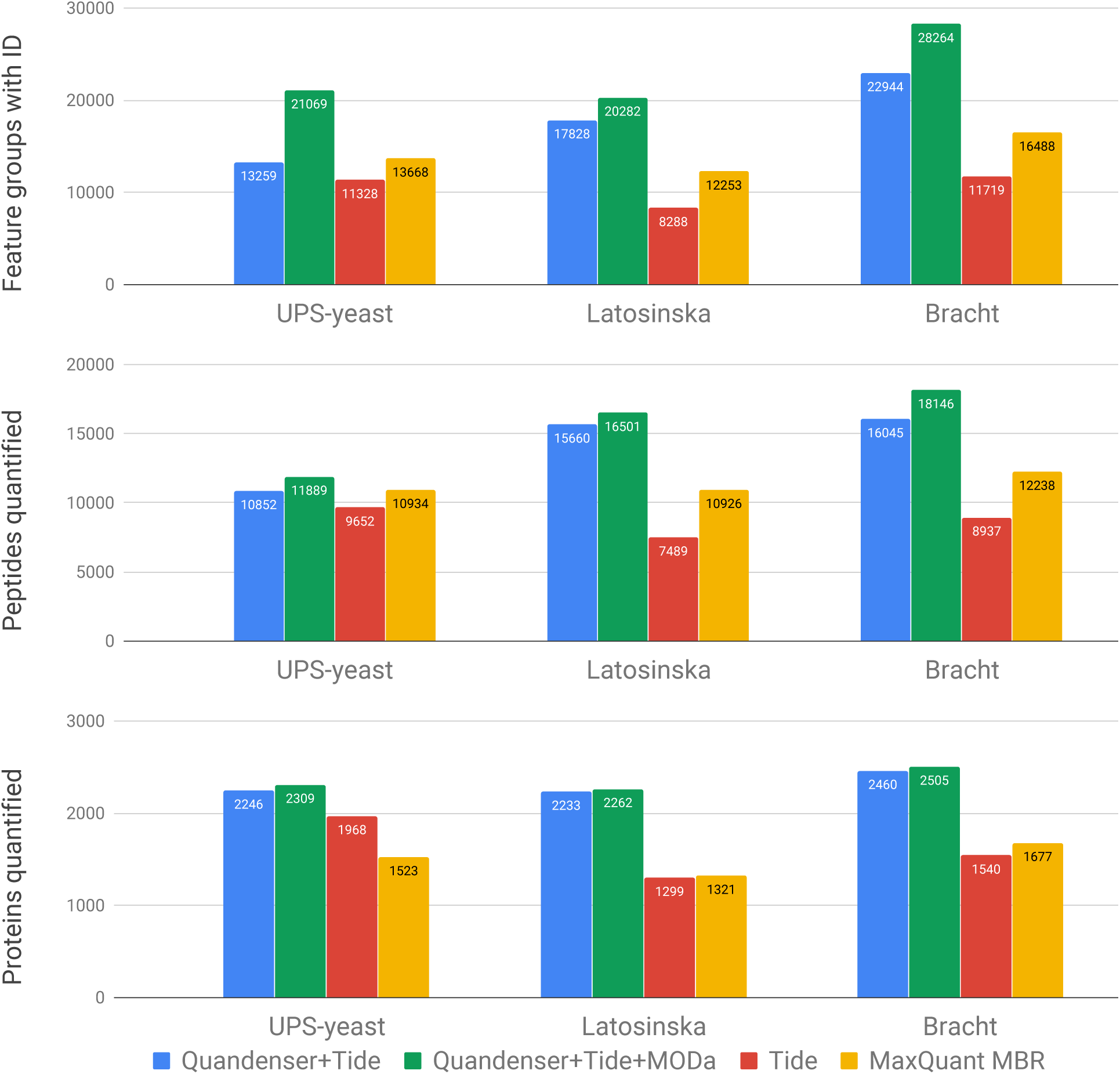
Using Quandenser, with and without the open modification search, increases the number of quantified peptides and proteins compared to not clustering and MaxQuant MBR. Performing an open modification search after a normal database search drastically increases the number of feature groups with identifications. We plot the number of feature groups with identification, quantified peptides and quantified proteins at 1% FDR after applying the maximum missing value criterion. The analyzed methods were Quandenser+Tide (**blue**), a cascade search of first Tide and subsequently MODa (**green**); Tide without Quandenser (**red**) and MaxQuant with MBR (**yellow**). For the comparison on protein level one should bear in mind that MaxQuant requires at least two unique peptides for protein identification.

Interestingly, searching with MODa without searching with Tide first actually decreased the sensitivity on the spiked-in proteins relative to only searching with Tide, even though more unique peptides were identified than by Tide. This is likely a result of the lower sensitivity of open modification searches on unmodified peptides, as a result of the increased search space. We indeed discovered several unmodified peptides from UPS proteins that were confidently identified by Tide but not picked up by MODa. In this engineered dataset the modified peptides did seem to follow the correct abundance pattern in the vast majority of the cases. In general, however, we should be careful about using modified peptides for quantification, as they are not guaranteed to follow the protein’s abundance pattern. On the other hand, quantifying modified peptides can be of great interest for understanding biological processes.

We also tested MSFragger with its large precursor tolerance (±500 Da) as an alternative open modification search engine. It produced more identifications than MODa, reducing the number of unidentified consensus spectra to around 40%, but also produced several dubious modifications. MSFragger could, therefore, be a good source for finding candidate peptide identifications, but some extra verification seems to be required for the moment.

To illustrate the utility of MS2 clustering, we used Novor on the 20 most frequently occurring unidentified spectra, followed by a BLAST search [25]. This interest was motivated by the fact that the identification rate for large clusters (> 80% for cluster size ≥ 16) was much higher than for small clusters (< 50% for cluster size ≤ 7). Using this approach, we found 2 distinct peptides from a capsid of a known yeast virus (UniProtKB: P32503 / GAG SCVLA) and another 2 distinct peptides from lysyl endopeptidase (UniProtKB: Q9HWK6 / LYSC PSEAE), the latter of which might have been used for improved protein digestion, although this was not mentioned in the original manuscript. All but one of these largest 20 unidentified spectrum clusters were identified as peptides from the 2 above-mentioned proteins or as modifications of already identified peptides of high-abundant proteins (Supplementary Table S2).

Furthermore, the benefit of having clustered on MS1 level allowed us to zoom in on feature groups without peptide identifications, but with the same abundance pattern as the UPS proteins. For this, we calculated the cosine distance between the expected abundance pattern and the observed abundance pattern, omitting missing values from the calculation, and selected the 200 feature groups with the smallest cosine distance (Figure 3, Supplementary Table S3). Of these 200 feature groups, 58 were identified through closer inspection as, often modified, UPS peptides and often came from chimeric spectra. One helpful approach in identifying these chimeric spectra was by filtering out the fragment peaks of an already identified peptide species and applying another open modification search [8].

**Figure 3:**
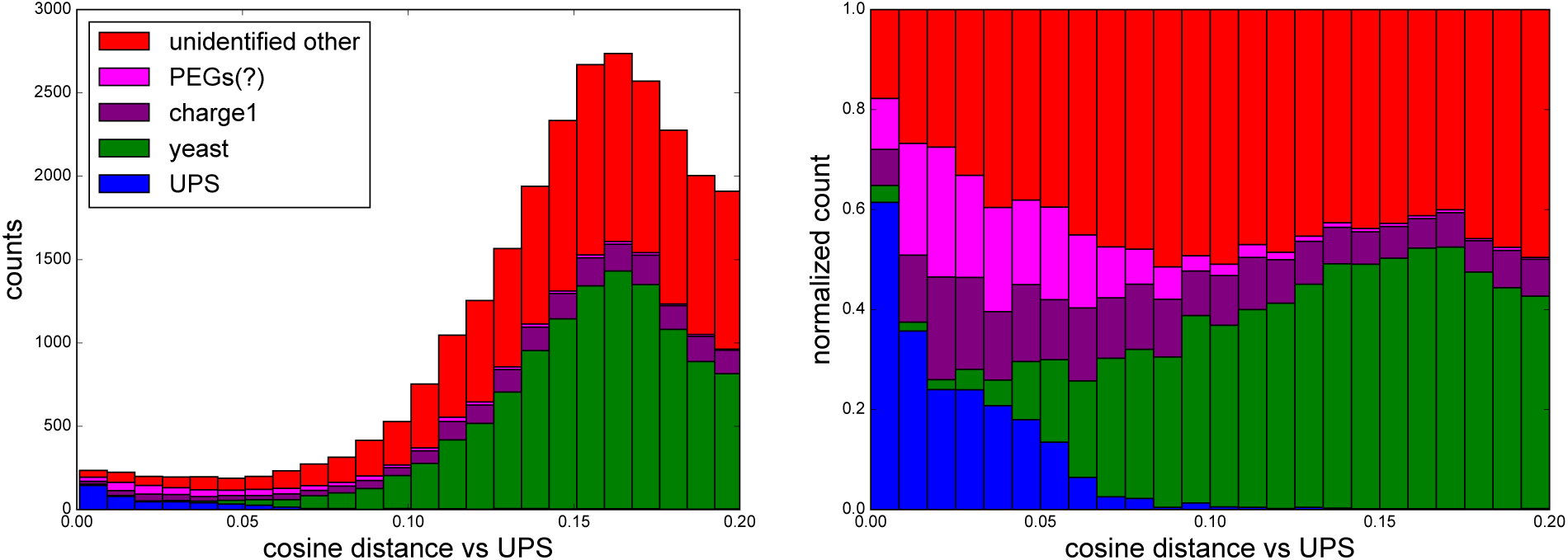
The number of feature groups and their origin as a function of dissimilarity to the UPS1 spike-in concentrations. The histogram displays the number (left pane) and relative number (right pane) of feature groups as a function of the cosine distance relative to the UPS1 spike-in concentrations. The vast majority of the identified feature groups had an abundance pattern that conformed to their origin. Still, a large proportion of feature groups remained unidentified, including a group that was tentatively identified as polyethylene glycols (PEGs), which exhibited an abundance pattern similar to the UPS1 proteins. The cosine distance between the yeast and UPS concentrations was 0.16 and we can indeed observe that the majority of the yeast peptides center around that value.

Interestingly, we also found 68 feature groups that had consensus spectra that contained ≥ 25 fragments between 100 and 200 Da. Based on the accurate masses of these fragments, these were carbohydrates or hydrocarbons and did not contain any nitrogen atoms (Supplementary Table S4). In total, we found 1 724 of these types of spectrum clusters, which mainly eluted towards the end of the runs. Their abundance pattern typically followed the UPS abundance pattern (Figure 3) and MS2 spectra associated with these feature groups generally remained unidentified. Based on their precursor mass differences and late retention times, these feature groups likely originated from polyethylene glycols (PEG), which might have been present as a contaminant in the UPS samples.

The putative identifications of a yeast virus, lysyl endopeptidase, and PEGs are examples that demonstrate that Quandenser gives its users the capability to identify unknown either abundant or differentially abundant compounds in their samples by targeting their spectra for identification. Possibly, these analytes could have been detected by other means, e.g. by using spectral libraries of known contaminants. However, for less engineered samples, Quandenser’s ability to help the user identifying unknown compounds can turn out to be indispensable.

Of the remaining 71 feature groups, 50 were from analytes with low precursor mass (< 1000 Da), mostly charge 1 ions, which are generally hard to identify. For 8 feature groups, the UPS abundance pattern was a result of deisotoping errors where isotopes of a UPS peptide were incorrectly counted towards the intensity of the feature. Finally, 13 feature groups remained unidentified and did not fit into any of the above categories, but usually had spectra that showed clear signs of chimericity or only had fragment ions spanning less than half of the peptide backbone making them hard to identify.

### Proteome mixture datasets

We processed all 3 proteome mixture (human-yeast, Shalit hela-ecoli, BoxCar hela-ecoli) datasets with Quandenser, followed by the aforementioned search engine strategies and protein quantification using Triqler (`--fold_change_eval=0.5`, just below the lowest spike-in ratio of log2(1.5) = 0.6). We consistently observed control of the differential abundance FDR and higher sensitivity relative to MaxQuant+Perseus and MaxQuant+EBRCT (Table 2, Supplementary Table S1) as well as reasonable estimates for the posterior distributions of fold changes (Supplementary Figure S9).

For the human-yeast mixture, for the 10 vs 5, 10 vs 3.3 and 5 vs 3.3 comparisons respectively, we found 24%, 46% and 85% more significant differentially abundant proteins with Quandenser+Tide+Triqler than in the MaxQuant+Perseus analysis of the original study (Supplementary Table S1). Compared to our own analysis with MaxQuant+Perseus, we obtained 13%, 16% and 175% more significant differentially abundant proteins. However, more importantly, in contrast to our pipeline, MaxQuant+Perseus and MaxQuant+EBRCT, again, did not control the differential abundance FDR, with respectively up to 10% and 21% observed differential abundance FDRs.

For the Shalit hela-ecoli set, no results were presented in the original study to which we could compare our results. Compared to our own analysis with MaxQuant+Perseus, we obtained an increase of at least 50% in the number of true positives in 4 out of 6 comparisons (Supplementary Table S1). The observed differential abundance FDRs for Quandenser+Tide+Triqler, MaxQuant+Perseus and MaxQuant+EBRCT were at most 6%, 6% and 13% respectively.

For the BoxCar hela-ecoli set, we used the MaxQuant+Perseus results deposited to PRIDE by the original authors. As the original study used a database that included Swiss-Prot and TrEMBL and we only used the Swiss-Prot part, the results are not directly comparable. Nevertheless, under the assumption that most protein groups from Swiss-Prot with TrEMBL correspond to a single SwissProt entry, Quandenser+Tide+Triqler obtained a comparable sensitivity for the BoxCar runs, but an increase of 35% for the Shotgun runs relative to the MaxQuant+Perseus results presented in the original study (Supplementary Table S1). Both methods managed to control the differential abundance FDR for the one-sided tests. For the two-sided tests, our pipeline controlled the differential abundance FDR for the BoxCar runs, but had an 18% observed differential abundance FDR for the Shotgun runs. However, the MaxQuant+Perseus analysis produced observed differential abundance FDRs of 71% and 50% respectively.

Due to the complex interplay of several parts of our pipeline it is hard to pinpoint exactly where the increased performance originates from. Nevertheless, we attempted to gain insights into the individual contributions of each of the steps by introducing several artificial filters on our identified feature groups that mimic the exclusion of certain parts of our pipeline. Specifically, we investigated the performance difference of including or excluding MS1 feature clustering, MS2 spectrum clustering and a peptide-level FDR threshold.

As a base case, we tested an approach that mimicked a typical input for a protein quantification method, that is, input on which no clustering was performed. For this, we evaluated the performance of our pipeline if we only used the feature groups that were (1) identified below 1% peptide-level FDR before MS2 clustering and (2) for which the number of missing values was ≥ *M* before MS1 clustering. In other words, we only retained feature groups for which at most *M* runs did not have an MS2 spectrum with the peptide identified. This approach is very similar to the pipeline Tide+Triqler without Quandenser/MBR from Table 2, with the addition of a peptide-level FDR threshold. We indeed observed a very similar performance of these two approaches (Supplementary Tables S1 and S33).

Second, we added to this set the feature groups for which at most *M* runs did not have an MS1 feature after applying MS1 clustering, still at 1% peptide-level FDR. The number of feature groups doubled, or even tripled, for all datasets through this addition, while maintaining high specificity (Supplementary Figure S13, Supplementary Table S33). This can mostly be attributed to the large reduction in missing values (Supplementary Figure S12) and is in line with what has previously been observed with several match-between-runs approaches [5, 46, 1].

Third, we included feature groups that had a peptide-level FDR below 1% after MS2 spectrum clustering. Although MS2 clustering has proven to be a valuable tool in increasing coverage of low abundant peptides [14], in our case, the MS1 clustering step already rescued these cases based on the MS1 features. Therefore, it was not very surprising that few feature groups were added in this step and that the addition of these feature groups had relatively little impact on sensitivity and specificity (Supplementary Figure S14, Supplementary Table S33). Nonetheless, MS2 spectrum clustering plays an important role in reducing the number of hypotheses and also is necessary for the intensity score filter introduced above. This filter relies on the MS2 spectrum clusters to select the most likely set of MS2 spectra for a given MS1 feature group.

Lastly, we investigated the influence of lower confident peptide identifications by adding the feature groups with a peptide-level FDR between 1% and 10%. This represented an appreciable number of feature groups and resulted in a modest increase in sensitivity (Supplementary Figure S15, Supplementary Table S33).

As has previously been observed, we found that many MS2 spectra remain unidentified and that a large number of MS1 feature groups remain without an assigned MS2 spectrum. The MS1 feature groups without identification at 1% peptide-level FDR constituted between 50% and 75% of all MS1 feature groups with MS2 spectrum, whereas adding MS1 feature groups without MS2 spectrum typically triples the number of feature groups (Supplementary Figure S16, Supplementary Table S33). As one could expect, the datasets with the highest proportions of unidentified MS2 spectra after the regular database search, also had the biggest increases in identifications by adding the open modification search results, bringing the identification rate across all datasets close to 50% (Supplementary Figure S5).

### Clinical datasets

The Latosinska dataset consisted of 413k MS2 spectra, resulting in 991k spectrum-feature matches after MS1 feature detection by Dinosaur. After filtering based on the intensity score, we were left with 83k feature groups, of which 47k had at least one spectrum cluster associated with them. This corresponded to 122k consensus spectra and 183k spectrum-feature matches to be searched, just 18% of the original number of spectrum-feature matches. This dataset contained a relatively large number of singleton clusters, i.e. clusters with only one spectrum. This highlights one of the benefits of doing quantification before identification in Quandenser. We can retain singleton clusters for which we can find MS1 features in multiple runs while discarding the singleton clusters for which this is not the case.

Searching the consensus spectra with Tide and/or MODa resulted in more identifications compared to MaxQuant on all levels (Figure 2). We are aware that the increase in quantified proteins can partly be explained by MaxLFQ’s requirement of having at least two peptides to quantify a protein. However, this is exactly the type of loss of sensitivity due to throwing evidence away prematurely in an attempt to artificially control the error rate that we try to address here. As we demonstrate in earlier work, Triqler can call proteins with only one peptide as differential abundant if the information from both the identification and quantification is very reliable.

Subsequent processing with Triqler (`--fold_change_eval=0.8`), resulted in more enriched functional annotation terms than applying Triqler directly on the MS2 search results (Figure 4, Supplementary Tables S5, S6, S7, S8, S9, S10, S11, S12, S13). There was a noticeable advantage for a cascaded Tide+MODa search, both in terms of the number of differentially abundant proteins, as well as in the number of enriched functional annotation terms. The original study found only a single protein at a 5% differential abundance FDR and 77 proteins with a *p* value below 0.05. These 77 proteins did not show any term enrichment in DAVID. The significant proteins called by MaxQuant+Perseus at the three FDR thresholds nor the 63 proteins with a *p* value below 0.05 showed any enriched terms either.

**Figure 4:**
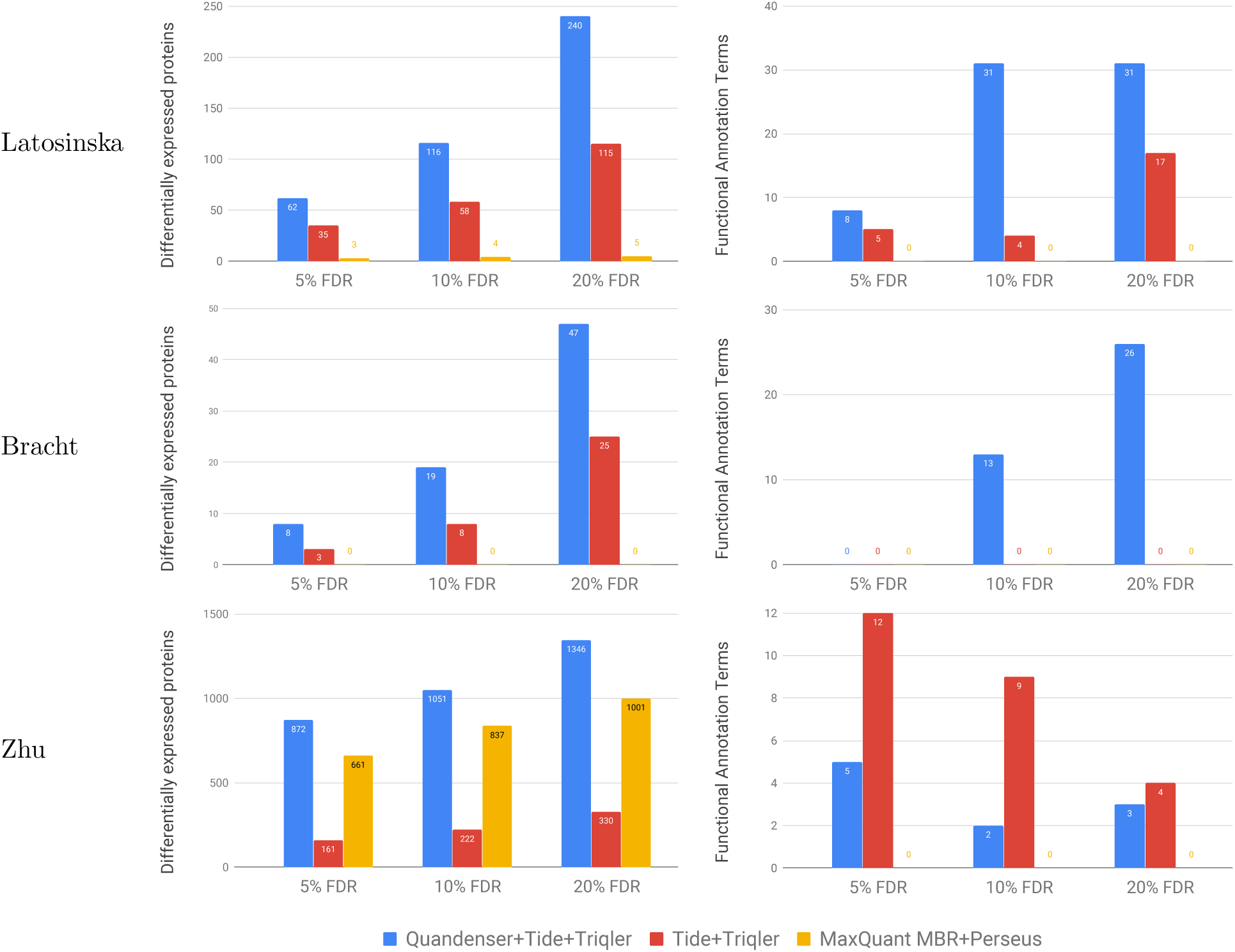
Benchmark of differentially abundant proteins and enriched functional annotation terms. The analyzed methods were our quantification-first approach, Quandenser+Triqler when using the search engine Tide (**blue**); and two identification-first approaches, Tide and Triqler without clustering nor a match-between-runs (MBR) feature (**red**) and MaxQuant with MBR followed by statistical analysis with Perseus (**yellow**). Overall, we discovered more differentially abundant proteins and enriched functional annotation terms with than without Quandenser. Notably, Quandenser+Triqler with Tide found enriched functional annotation terms for the Bracht set for which no enrichments were previously found. The left plots show the number of differentially abundant proteins at 3 differential abundance FDR thresholds. The plots on the right show the number of significant functional annotation terms we discovered with DAVID using the sets obtained in the left plots. Note that the FDR reported in the plots on the right refer to the differential abundance FDR and not the functional annotation term FDR, which was kept fixed at 5%.

The Bracht dataset contained 1.01M MS2 spectra, which were assigned a total of 1.47M spectrum-feature matches after feature detection. Processing with Quandenser resulted in 69k feature groups in total and 45k with at least one spectrum cluster. This left 106k consensus spectra, with 150k spectrum-feature matches to be searched, which was only 10% of the original number of spectrum-feature matches.

Again, we observed an increase in the number of identifications on all levels compared to MaxQuant (Figure 2). Analysis of the Tide search results with Triqler (`--fold_change_eval=0.5`; the fold change threshold used in the original study was log2(1.5) = 0.58) resulted in multiple differentially abundant proteins for all three searches, even at as low as 5% FDR (Figure 4). Conversely, neither the original study nor MaxQuant+Perseus found any differentially abundant proteins at 5% FDR.

In the original paper, 7 proteins with a *p* value below 0.05 and high fold change differences were subjected to verification through gene expression analysis, as well as targeted analysis with MRM. Notably, the 4 proteins that showed a consistent relationship with increasing fibrosis stages in both experiments (FBLN5, LUM, COL14A1, and MFAP4) were all discovered at 10% FDR, as was one protein that showed significance in the gene expression analysis but only partial statistical significance in the MRM analysis (TAGLN). The 2 proteins with the least consistent relation of expression levels with increasing fibrosis stages (CSRP2 and CNN2) were not discovered at 10% FDR, though CSRP2 was called significant at 20% FDR. In the original study, these 2 proteins actually obtained a lower *p* value than the other 5 proteins, and Quandenser+Triqler, thus, seemed to give a better ordering of these 7 proteins.

Moreover, functional annotation analysis with DAVID actually resulted in several significant terms for the 10% and 20% term-FDR thresholds (Supplementary Table S14, S15, S16, S17, S18, S19, S20, S21, S22). Compellingly, several terms related to the extracellular matrix were found significant, which was also pinpointed as a entity of interest in the original paper based on earlier studies.

Finally, we wanted to test the applicability of the method to the Zhu dataset, which contained samples of a small number (10–100) of cells. This dataset comprised 593k MS2 spectra, assigned to 1.02M spectrum-feature matches. Processing with Quandenser resulted in 73k feature groups of which 52k had at least one spectrum cluster. This left 117k consensus spectra, corresponding to 187k spectrum-feature matches to be searched, 18% of the original number. We set the log2 fold change threshold `--fold_change_eval=1.0` in Triqler, corresponding to the threshold employed in the original study.

The number of identified feature groups and unique peptides were closer between Quandenser and MaxQuant for this particular set (Supplementary Figure S6). Nevertheless, Quandenser+Triqler managed to discover more significant proteins across all tested FDR thresholds compared to MaxQuant+Perseus (Figure 4). Using Tide as a search engine, we found 703 significant proteins at 2% FDR, considerably more than the 304 proteins at the same FDR found in the MaxQuant+Perseus analysis of the original study. Again, we found enriched functional annotation terms associated to several sets of significant proteins using Quandenser+Triqler (Supplementary Table S23, S24, S25, S26, S27, S28, S29, S30, S31). No enriched terms could be found for MaxQuant+Perseus analysis, neither from the original study nor from our own reanalysis. Interestingly, using Triqler without Quandenser appeared more sensitive in the functional annotation enrichment analysis, as did using stricter FDR thresholds.

## Discussion

Here, we demonstrated the utility of a quantification-first approach in which we cluster both MS1 features and MS2 spectra prior to identifying spectra, peptides, and proteins of interest. While the idea to quantify features without an explicit identity is well explored in the literature [44, 31, 9], we here have demonstrated the idea’s usefulness in combination with clustering. Not only does this approach provide several new ways of obtaining unidentified analytes of interest, but by combining Quandenser with Triqler, we discovered substantially more differentially abundant proteins and enriched functional annotation terms than MaxQuant+Perseus.

We demonstrated that clustering was effective in increasing the coverage of the examined samples. By first reduced sets of spectra, we could apply relatively computationally expensive search strategies. First, an open modification search increased the number of identified consensus spectra by 47% for the UPS-yeast dataset and 14 − 20% for the clinical datasets. Second, *de novo* searches on large MS2 spectrum clusters of the UPS-yeast dataset resulted in the identification of peptides and proteins not present in the database. Third, the similarity of feature groups’ abundance patterns across runs indicate that they originated from the same group of proteins. By restricting the search to the 48 UPS proteins we found that many analytes with a similar abundance to the spiked-in proteins were modified UPS peptides. Unfortunately, such targeted searches easily cause false reinforcement of protein quantities, but it does reveal peptides and modifications we are missing out on. This approach also revealed a group of analytes that covaried with the UPS proteins, likely to be polyethylene glycols. These analytes would not have come to our attention in a traditional identification-first approach. Our intention was not to demonstrate that the modification and *de novo* searches we applied here are the best way of achieving increased proteome coverage. Rather, our analysis presents a first look into the part of the proteome that normally is ignored, with hopefully many more discoveries yet to come.

Another benefit of combining clustering on MS1 with MS2 level is that we can include quantification information in the selection of MS2 spectrum clusters of interest. This addresses a frequently observed phenomenon in MS2 spectrum clustering in which the majority of the clusters only contain a single spectrum, known as singleton clusters [10, 12, 39]. These could be spurious MS2 fragmentation events, which should preferably not be matched by a search engine. On the other hand, these could be low abundant peptides rarely selected for fragmentation. The Quandenser workflow can separate these cases by only retaining fragment spectra from analytes that were quantified across several runs. Furthermore, by using the agreement of the MS2 spectra within an MS2 spectrum cluster regarding which MS1 feature group was most likely targeted, we managed to set up the efficient intensity score filter, which drastically reduced the number of hypotheses with few false negatives.

The clustering steps further unveil the potential of integrated quantification and identification error models such as Triqler. Particularly, using matches-between-runs with feature-feature match error rates contributes to controlling the differential abundance FDR. Notably, even though the number of peptide and protein identifications favors our method, we want to emphasize that it is quantitative reliability that is key in the end. By using statistical models with error propagation instead of sets of (arbitrary) thresholds, our pipeline consistently achieved control of the differential abundance FDR, whereas both MaxQuant+Perseus and MaxQuant+EBRCT markedly failed to do so.

The functional enrichment term analysis by DAVID showed promising results on the clinical datasets, finding enriched terms for all three sets where MaxQuant+Perseus repeatedly showed no term enrichments. However, care should be taken in interpreting these results as the methods used here might introduce biases that analysis tools such as DAVID are sensitive to. For example, our method allows the quantification of proteins with only a single confident peptide in which the prior still has a strong influence. These proteins are far less likely to reach the differential abundance significance threshold but are still included in the background of identified proteins. Ideally, one would feed the posterior distributions directly into a downstream (Bayesian) method, without setting thresholds on differential abundance significance. With the recent rise of Bayesian methods for protein quantification, we expect such methods to become a topic of interest in the near future.

Several improvements could still be made to increase the sensitivity of Quandenser and the quantification-first pipeline in general. As with other label-free quantification approaches, Quandenser is highly dependent on the ability to reliably extract features from the MS1 chromatograms, which becomes harder as the density of MS1 features increases [2, 32]. However, Quandenser could easily be extended to fractionated data, or even to ion-mobility data [7] using the same clustering principles as employed here. Moreover, Quandenser forms consensus spectra by a weighted averaging of the peak intensities of the spectra in a cluster [10, 39]. However, the search engine scores of consensus spectra are seldom higher than the best scoring constituent spectra. Improving the process to form consensus spectra would have benefits for other types of analysis, such as the formation of spectral libraries. Another problem with the current methods for clustering MS2 spectra is the so-called *chimeric spectra*, i.e. spectra containing product-ions from multiple peptides, that can contaminate clusters [39]. Solving how to cluster partial fragment spectra based on similarities of subseries of the full spectrum would overcome this problem and open up avenues for applying Quandenser-like processing of data from data independent analysis (DIA) mass spectrometry.

Finally, we specifically note the great potential of the quantification-first approach for processing datasets larger in sample size. First, by only requiring quantification to take place once, we remove one of the most time-consuming parts of re-analysis with different search parameters or engines. Second, by reducing the number of spectra that we need to analyze, we reduce the search time as well as lower the number of hypotheses we need to test when analyzing the subsequently matched spectra. The combination of these two techniques allows a wide variety of, potentially computationally expensive, search strategies to be applied to the quantification data. Especially with regards to the constant stream of new search engines, this provides an easy way for users to efficiently interrogate their data.

The quantification-first approach in label-free protein quantification, thus, provides an attractive alternative to the traditional identification-first approach. Through the use of unsupervised clustering, we condensed the data into a comprehensive format that retained the relevant information and thereby allow the researcher to spend more time on a reduced set of hypotheses. By subsequently propagating the feature-level error rates to probabilistic protein quantification methods, the bounds of sensitivity and specificity in LFQ are extended considerably. Already, we can see the benefits of this approach in terms of coverage and sensitivity using the techniques presented here, but many more modes of interpretation are available, ready to be applied.

## Acknowledgments

This work was supported by a grant from the Swedish Research Council (grant 2017-04030). We would like to thank Timothy Bergström, Kungliga Tekniska Högskolan, for writing the Python converter for combining MS1 spectra in the BoxCar runs.

## Conflict of interest

The authors declare that they have no conflict of interest.

