## Supplementary Information for "Focus on the spectra that matter by clustering of quantification data in shotgun proteomics"

March 18, 2020

#### 1 Updated graphical model for Triqler

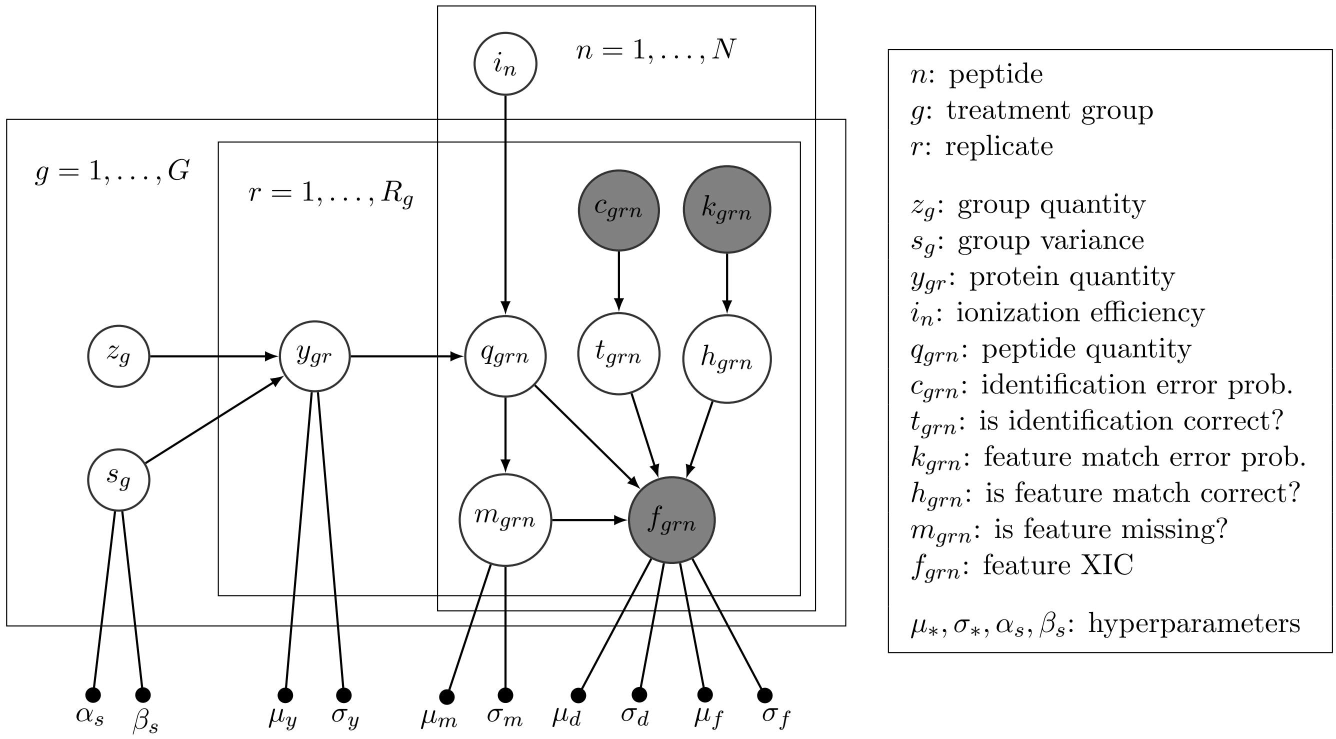


Figure S1: **Updated probabilistic graphical model for Triqler with $k_{grn}$ and $h_{grn}$ nodes for feature-feature match PEPs.** The protein has $N$ peptides, $G$ treatment groups and $R_g$ replicates per treatment group. Gray nodes denote observed variables, whereas white nodes denote latent variables. A concrete example is presented in section 1.1.

For the new indicator variable $h_{grn}$ we define:

$$h_{grn} = \begin{cases} 0, & \text{if feature match is correct} \\ 1, & \text{if feature match is incorrect} \end{cases}$$

The term $p(f_{grn} \mid q_{grn}, t_{grn}, m_{grn})$ is now replaced by $p(f_{grn} \mid q_{grn}, t_{grn}, m_{grn}, h_{grn})$, which is defined by:

$$p(f_{grn} = \text{NaN} \mid m_{grn} = 0, t_{grn} = 0) = 0$$
$$p(f_{grn} = \text{NaN} \mid q_{grn} = a, m_{grn} = 1, t_{grn} = 0, h_{grn} = 0) = 1$$
$$p(f_{grn} = \text{NaN} \mid h_{grn} = 1) = p(f_{grn} = \text{NaN} \mid t_{grn} = 1, h_{grn} = 0) = \text{sigm}(w; \mu_m, \sigma_m)$$
$$p(f_{grn} = x \mid m_{grn} = 1, t_{grn} = 0) = 0$$
$$p(f_{grn} = x \mid q_{grn} = a, m_{grn} = 0, t_{grn} = 0, h_{grn} = 0) = \text{hypsec}(a - x; \mu_d, \sigma_d)$$
$$p(f_{grn} = x \mid h_{grn} = 1) = p(f_{grn} = x \mid t_{grn} = 1, h_{grn} = 0) = (1 - \text{sigm}(w; \mu_m, \sigma_m)) \cdot \text{hypsec}(w - x; \mu_d, \sigma_d)$$

with $w = \left(\prod_W f_{g'r'n}\right)^{1/|W|}$ and $W = \{(g', r') \mid (g', r') \neq (g, r), f_{g'r'n} \neq \text{NaN}\}$, the geometric mean of all non-missing other features from the same feature group.

For the term $p(h_{grn} \mid k_{grn})$ we simply have

$$p(h_{grn} = 1 \mid k_{grn}) = k_{grn}$$
$$p(h_{grn} = 0 \mid k_{grn}) = 1 - k_{grn}$$

#### 1.1 Example

As an example, let us consider one of the UPS proteins, P08263ups|GSTA1_HUMAN_UPS, from the UPS-Yeast mix dataset. This protein has 4 peptide identifications and, ideally, we want to show that this protein is present in different concentrations in each of the 3 groups, where the UPS proteins were spiked in at 5, 10 and 25 fmol.

For each combination of sample and peptide, we have 3 pieces of information: the feature intensity (XIC, $f_{grn}$), the posterior probability of the MS2 cluster identification to be incorrect ($P_{id}$, $c_{grn}$) and the posterior probability of the MS1 feature to be incorrectly matched to one of the MS1 features with an MS2 cluster identification ($P_{match}$, $k_{grn}$):

| group | 25 fmol ($g = 1$) | | | | | | | | |
|---|---|---|---|---|---|---|---|---|---|
| P08263ups\|GSTA1_HUMAN_UPS | C-25fmol-R1 ($r = 1$) | | | C-25fmol-R2 ($r = 2$) | | | C-25fmol-R3 ($r = 3$) | | |
| | XIC $f_{11n}$ | $P_{id}$ $c_{11n}$ | $P_{match}$ $k_{11n}$ | XIC $f_{12n}$ | $P_{id}$ $c_{12n}$ | $P_m$ $k_{12n}$ | XIC $f_{13n}$ | $P_{id}$ $c_{13n}$ | $P_m$ $k_{13n}$ |
| ISNLPTVK ($n = 1$) | 61.9 | 0.004 | 0.027 | 73.0 | 0.003 | 0 | NaN | 0.004 | 0.066 |
| YFPAFEK ($n = 2$) | 103.5 | 0.02 | 0 | 92.2 | 0.02 | 0 | 95.1 | 0.02 | 0 |
| KFLQPGSPRKPPMDEK ($n = 3$) | 0.12 | 0.93 | 0.04 | 0.20 | 0.93 | 0.02 | 0.24 | 0.93 | 0.07 |
| FLQPGSPR ($n = 4$) | 66.3 | 0.028 | 0 | 55.4 | 0.028 | 0 | 84.3 | 0.028 | 0 |

| group | 10 fmol ($g = 2$) | | | | | | | | |
|---|---|---|---|---|---|---|---|---|---|
| P08263ups\|GSTA1_HUMAN_UPS | D-10fmol-R1 ($r = 1$) | | | D-10fmol-R2 ($r = 2$) | | | D-10fmol-R3 ($r = 3$) | | |
| | XIC $f_{21n}$ | $P_{id}$ $c_{21n}$ | $P_{match}$ $k_{21n}$ | XIC $f_{22n}$ | $P_{id}$ $c_{22n}$ | $P_m$ $k_{22n}$ | XIC $f_{23n}$ | $P_{id}$ $c_{23n}$ | $P_m$ $k_{23n}$ |
| ISNLPTVK ($n = 1$) | 7.56 | 0.004 | 0.049 | 20.6 | 0.004 | 0 | 18.0 | 0.004 | 0.054 |
| YFPAFEK ($n = 2$) | NaN | 0.02 | 0.017 | 22.0 | 0.02 | 0.012 | 26.6 | 0.02 | 0 |
| KFLQPGSPRKPPMDEK ($n = 3$) | 0.23 | 0.93 | 0.020 | 0.13 | 0.93 | 0.033 | 0.11 | 0.93 | 0.015 |
| FLQPGSPR ($n = 4$) | 16.7 | 0.028 | 0.15 | 15.0 | 0.028 | 0.12 | 19.53 | 0.028 | 0.20 |

| group | 5 fmol ($g = 3$) | | | | | | | | |
|---|---|---|---|---|---|---|---|---|---|
| P08263ups\|GSTA1_HUMAN_UPS | E-5fmol-R1 ($r = 1$) | | | E-5fmol-R2 ($r = 2$) | | | E-5fmol-R3 ($r = 3$) | | |
| | XIC $f_{31n}$ | $P_{id}$ $c_{31n}$ | $P_{match}$ $k_{31n}$ | XIC $f_{32n}$ | $P_{id}$ $c_{32n}$ | $P_m$ $k_{32n}$ | XIC $f_{33n}$ | $P_{id}$ $c_{33n}$ | $P_m$ $k_{33n}$ |
| ISNLPTVK ($n = 1$) | 6.96 | 0.004 | 0.066 | 7.27 | 0.004 | 0 | 2.38 | 0.004 | 0.009 |
| YFPAFEK ($n = 2$) | 4.81 | 0.02 | 0.017 | 1.83 | 0.02 | 0.010 | 8.68 | 0.02 | 0.007 |
| KFLQPGSPRKPPMDEK ($n = 3$) | 0.14 | 0.93 | 0 | 0.39 | 0.93 | 0 | 0.13 | 0.93 | 0.009 |
| FLQPGSPR ($n = 4$) | 10.9 | 0.028 | 0.09 | 5.36 | 0.028 | 0.077 | 10.9 | 0.028 | 0.055 |

We can plot the posterior distributions on several levels for this particular protein using some extra functionality of the Triqler python package:

```
python -m triqler.distribution.plot_posteriors --protein_id P08263ups
    --fold_change_eval 0.8 --decoy_pattern decoy_ --max_plot_fold_change 4.0
    --spike_in_concentrations 25,10,5
    UPS_Yeast_mix.quant_rows.tide_concat_peptides.tsv
```

On protein level (Figure S2), we can see the influence of the missing values in C-25fmol-R3 (first pane, red series) and D-10fmol-R1 (second pane, blue series). These missing values shift some of the probability to lower relative protein quant levels, but due to the strong evidence from the other peptides, this shift is only modest. Also, note that these posteriors are not necessarily normal distributions.

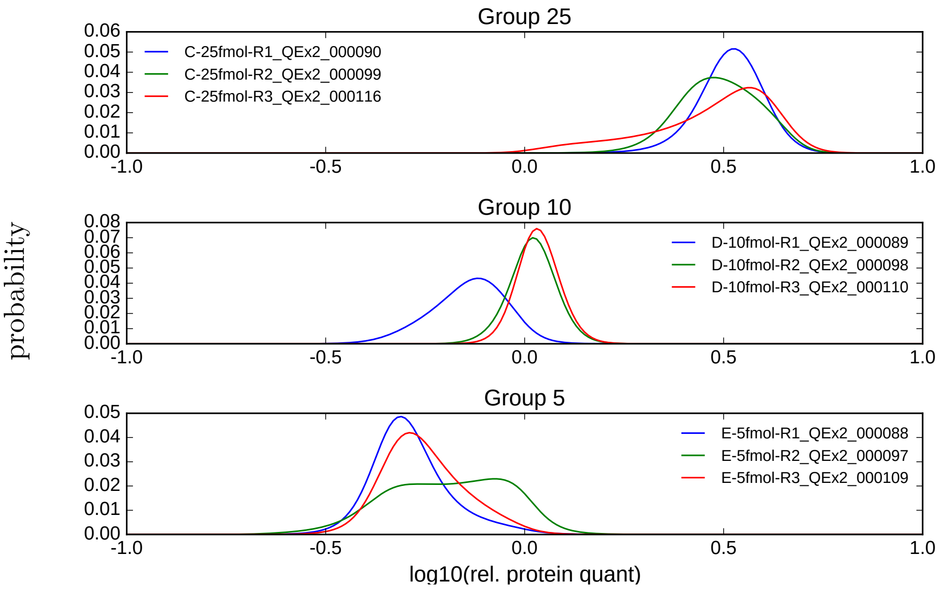


Figure S2: **Posterior distributions for protein abundances as calculated by Triqler.** We plot the probabilities as a function of the $\log_{10}$ relative protein abundance for each of the 9 samples, grouped by spike-in concentration. We can see some clear contributions of the individual peptides, such as missing values (Group 10, blue series) and disagreement between different peptides (Group 5, green series).

On experimental group level (Figure S3), we can still see traces of the contributions of the individual proteins, especially in the different widths of the group distributions. However, due to the averaging over the three replicates, we end up with distributions that are very close to normal distributions.

On fold change level (Figure S4, note that the abundance scale changed from $\log_{10}$ to $\log_2$ compared to the treatment group posterior plots), the width of the distributions for 25 vs 10 is noticably smaller than for the other two comparisons, which can be attributed to the larger uncertainty in abundance for the 5 fmol protein and group distributions.

Some other notable features of the graphical model:

- For the samples that have an MS2 spectrum in the MS2 spectrum cluster that identified the peptide in question, the feature match error probability $P_m$ is 0, e.g. $k_{121}$. This neglects the possibility that the MS2 spectrum was erroneously assigned to the MS1 feature group or that the MS2 spectrum was erroneously assigned to the MS2 spectrum cluster. We assume these probabilities to be small, but incorporating these in an extension of the model might be addressed in future work.

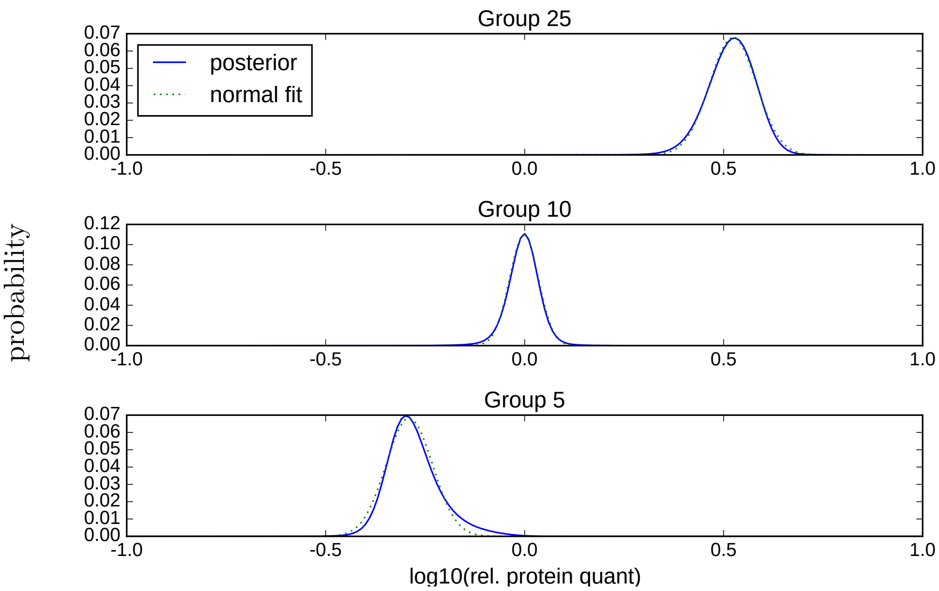


Figure S3: **Posterior distributions for experimental group mean abundances as calculated by Triqler.** We plot the probabilities as a function of the $\log_{10}$ experimental group mean abundance for each of the 3 spike-in concentrations. Group 10 clearly has a narrower distribution, owing to the better agreement between peptides and replicates. For illustration puposes, we have included least square fittings of normal distributions to the calulated posterior distributions.

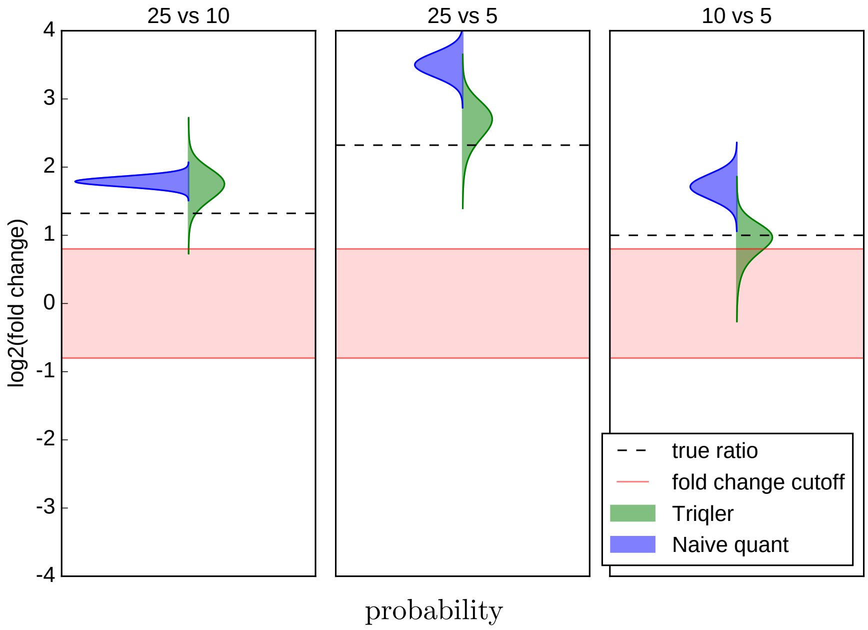


Figure S4: **Posterior distributions for fold changes between experimental groups as calculated by Triqler (green) and a naive quantification method (blue).** We plot the probabilities as a function of the $\log_2$ fold change between experimental groups. The naive quantification method summed up the 3 most intense peptides and imputed missing values by the row mean. We plot the naive quantification method's results as normal distributions using the mean and standard error of the mean. Triqler manages to produce posterior distributions that are closer to the true ratios.

- Conversely, a non-zero $P_m$ value, e.g. $k_{111}$, means that the sample in question did not have an MS2 spectrum that could be associated to the peptide but did have an MS1 feature that could be matched to one of the MS1 features in another sample in the minimum spanning tree alignment.
- Usually, only one MS2 spectrum cluster will yield a confident peptide identification for an MS1 feature group. In this example, we can see this for YFPAFEK, KFLQPGSPRKPPMDEK and FLQPGSPR, where the identification probability is the same across all samples. In the case of ISNLPTVK, there were 2 MS2 spectrum clusters with the same peptide identification, one with $P_{id} = 0.003$ and one with $P_{id} = 0.004$. This happens if the MS2 spectra are not sufficiently similar for MaRaCluster to cluster them together, for example due to chimericity.
- The MS2 spectrum cluster for KFLQPGSPRKPPMDEK does not have a reliable peptide identification ($P_{id} = 0.93$), and therefore its contribution to the protein quantification will be penalized.
- Both $f_{131}$ and $f_{212}$ are missing values. Combined with the high XIC of $f_{111}$ and $f_{121}$, the model will, for example, infer that $f_{131}$ is missing completely at random.
- The feature match probability $k_{234}$ for the MS1 feature with XIC 19.53 is rather high. This will penalize the contribution of $f_{234}$, but as it actually agrees with the other confidently identified peptides its contribution is still important to .

#### 2 Identification statistics

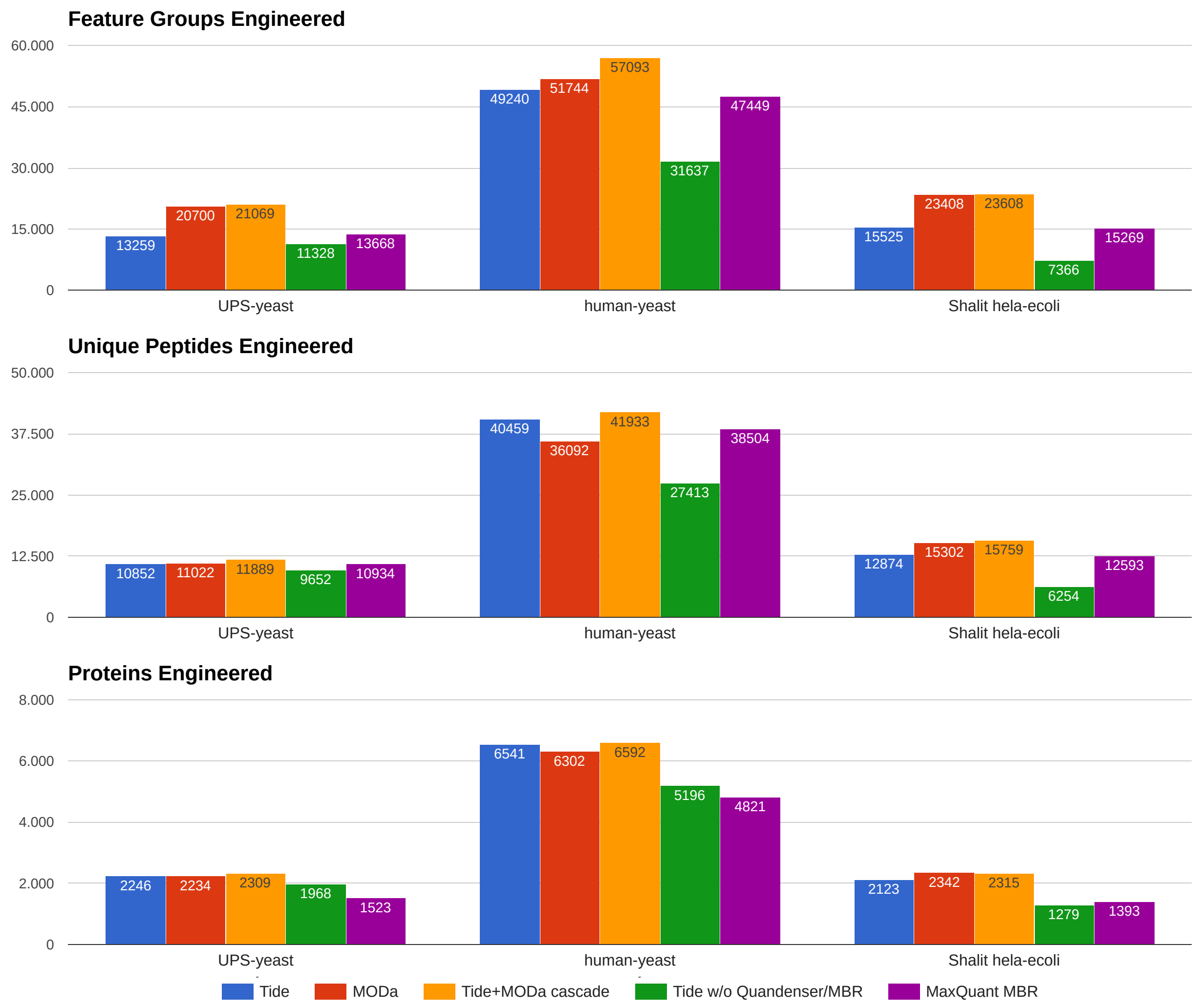


Figure S5: **Using a cascaded search with an open modification search as second search increases the number of identifications on all levels on three engineered datasets.** The analyzed methods were Quandenser+Triqler when using the search engine Tide (**blue**), MODa (**red**), and a cascade search of first Tide and subsequently MODa (**yellow**); Tide and Triqler without MaRaCluster and the matches-between-runs (MBR) feature (**green**) and MaxQuant with MBR followed by statistical analysis with Perseus (**orange**). A comparison of the number of feature groups with a peptide identification, unique peptides and proteins at 1% identification FDR shows the superior performance of using a cascaded search which includes an open modification search through MODa. It should be noted that MaxQuant requires at least two unique peptides for protein identification and therefore the difference in the number of identifications on protein level cannot directly be compared.

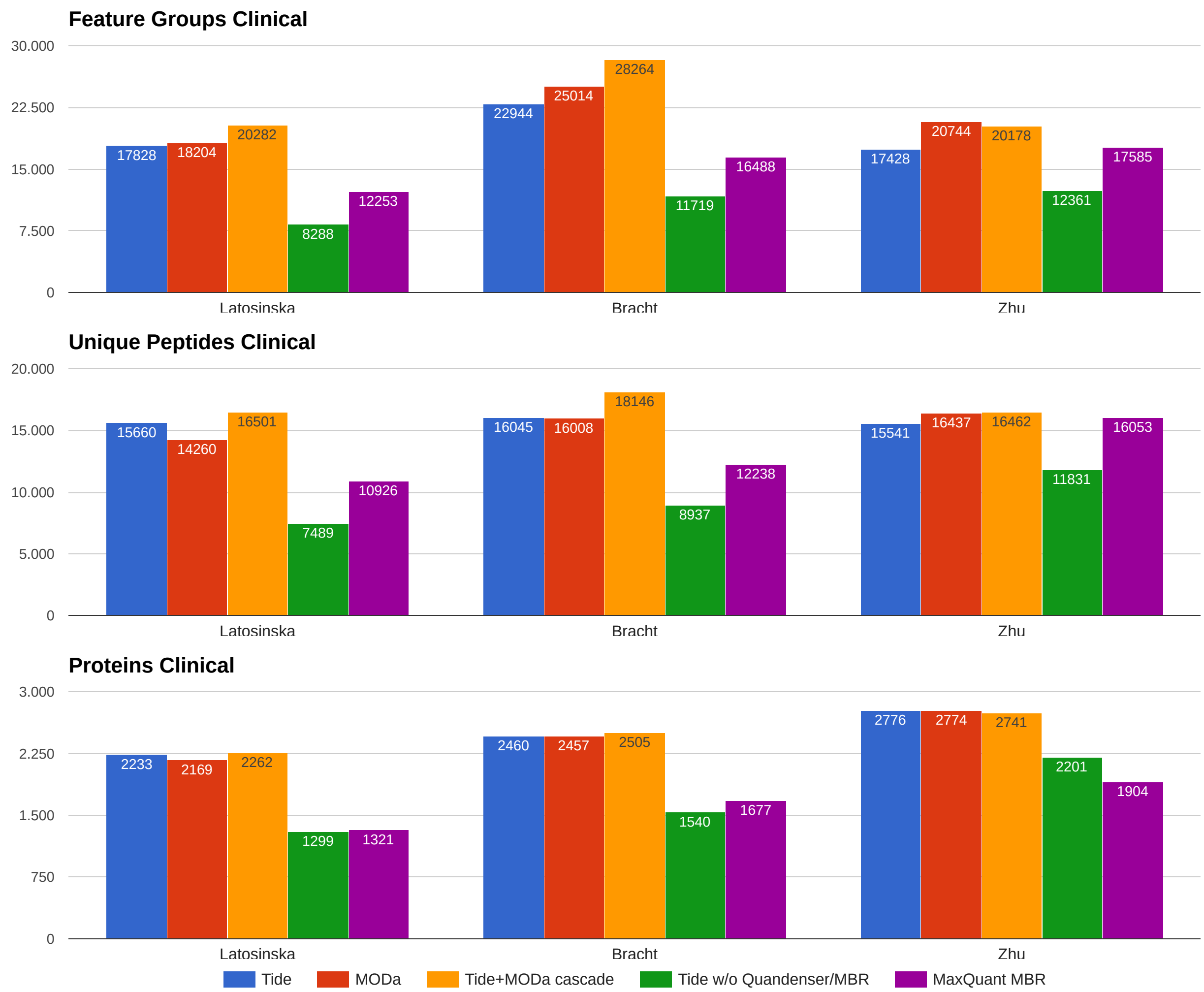


Figure S6: **Using a cascaded search with an open modification search as second search increases the number of identifications on feature group and peptide levels for three clinical datasets.** The analyzed methods were Quandenser+Triqler when using the search engine Tide (**blue**), MODa (**red**), and a cascade search of first Tide and subsequently MODa (**yellow**); Tide and Triqler without MaRaCluster and the matches-between-runs (MBR) feature (**green**) and MaxQuant with MBR followed by statistical analysis with Perseus (**orange**). A comparison of the number of feature groups with a peptide identification, unique peptides and proteins at 1% identification FDR shows the superior performance of using a cascaded search which includes an open modification search through MODa. It should be noted that MaxQuant requires at least two unique peptides for protein identification and therefore the difference in the number of identifications on protein level cannot directly be compared.

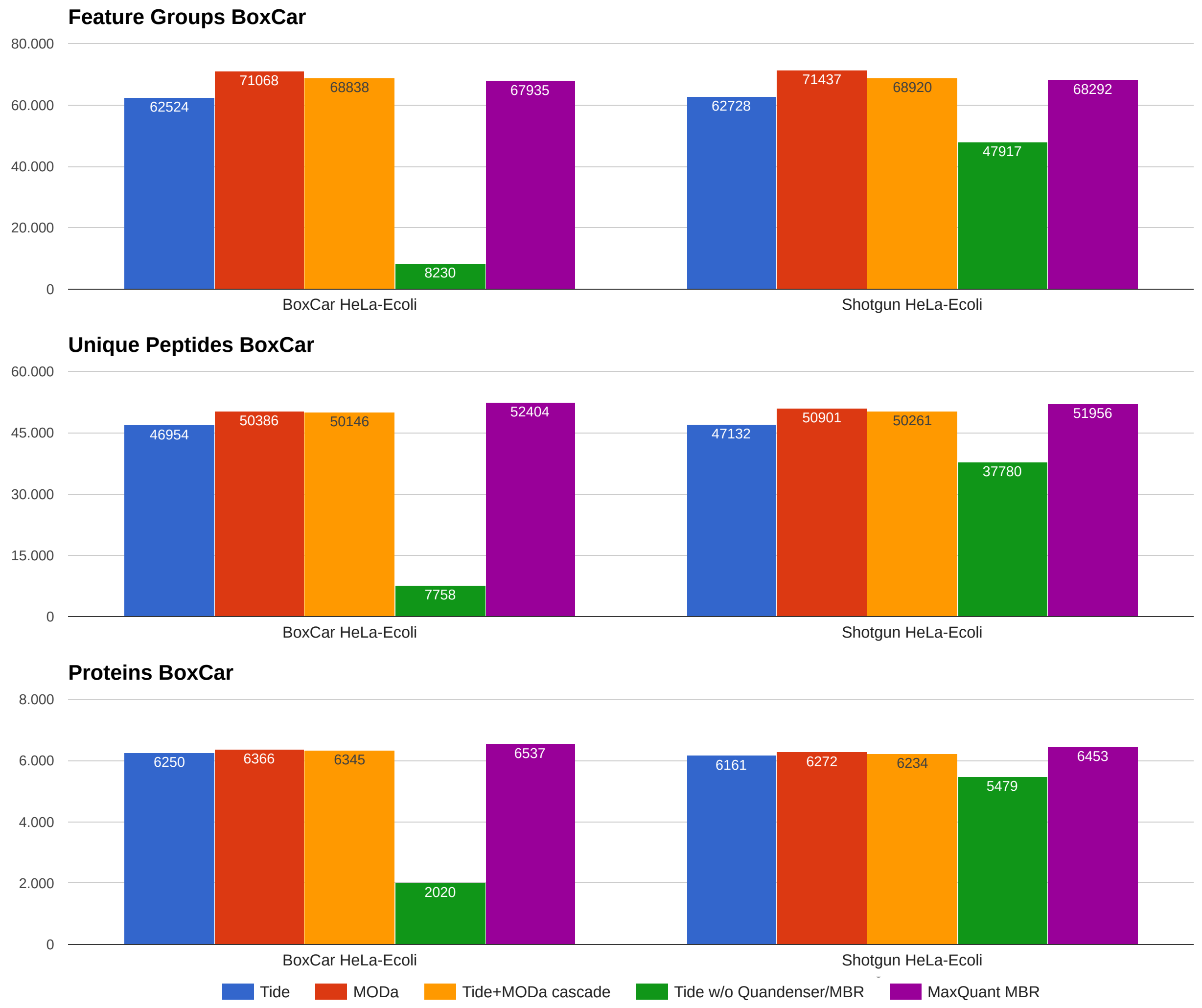


Figure S7: **Using a cascaded search with an open modification search as second search increases the number of identifications on all levels for both the Shotgun and BoxCar runs from the BoxCar manuscript.** The analyzed methods were Quandenser+Triqler when using the search engine Tide (**blue**), MODa (**red**), and a cascade search of first Tide and subsequently MODa (**yellow**); Tide and Triqler without MaRaCluster and the matches-between-runs (MBR) feature (**green**) and MaxQuant with MBR followed by statistical analysis with Perseus (**orange**). A comparison of the number of feature groups with a peptide identification, unique peptides and proteins at 1% identification FDR shows the superior performance of using a cascaded search which includes an open modification search through MODa. It should be noted that we used the MaxQuant output files from the original BoxCar manuscript, in which the database contained all proteoforms in Swiss-Prot and TrEMBL, whereas we only searched against the proteins in Swiss-Prot. Therefore, the difference in the number of identifications between MaxQuant and Quandenser+Triqler cannot directly be compared. Also, since the BoxCar approach only samples very few MS2 spectra, peptide identifications need to be inferred using MBR with respect to the Shotgun runs. This explains the poor behavior of Tide without Quandenser/MBR on the BoxCar runs and its decent behavior on the Shotgun runs.

#### 3 Differential expression statistics

**UPS-yeast**

| UPS1 spike-in concentration [fmol] | 25 vs 10 | | 25 vs 5 | | 10 vs 5 | |
|---|---|---|---|---|---|---|
| Method | tp | fp | tp | fp | tp | fp |
| **Quandenser+Tide+Triqler** | **43** | **1** | **45** | **1** | **35** | **0** |
| MaxQuant MBR+Perseus | | | | | | |
| - $S_0 = 0.0$ | 37 | 42 | 35 | 124 | 24 | 7 |
| - $S_0 = 0.3$ | 39 | 9 | 42 | 15 | 36 | 1 |
| - $S_0 = 0.7$ | 34 | 9 | 40 | 6 | 24 | 1 |
| - $S_0 = 1.0$ | 28 | 2 | 41 | 5 | 1 | 0 |
| MaxQuant MBR+EBRCT | 43 | 1 | 40 | 9 | 0 | 0 |

**Shalit hela-ecoli**

| E. coli spike-in concentration [ng] | 3 vs 7.5 | | 3 vs 10 | | 3 vs 15 | | 7.5 vs 10 | | 7.5 vs 15 | | 10 vs 15 | |
|---|---|---|---|---|---|---|---|---|---|---|---|---|
| Method | tp | fp | tp | fp | tp | fp | tp | fp | tp | fp | tp | fp |
| **Quandenser+Tide+Triqler** | **190** | **7** | **198** | **4** | **229** | **0** | **0** | **0** | **194** | **3** | **138** | **0** |
| MaxQuant MBR+Perseus | | | | | | | | | | | | |
| - $S_0 = 0.0$ | 75 | 5 | 96 | 38 | 129 | 246 | 0 | 1 | 100 | 27 | 41 | 2 |
| - $S_0 = 0.3$ | 126 | 7 | 134 | 6 | 137 | 9 | 0 | 0 | 128 | 4 | 125 | 7 |
| - $S_0 = 0.7$ | 117 | 7 | 134 | 4 | 137 | 7 | 0 | 0 | 125 | 6 | 4 | 0 |
| - $S_0 = 1.0$ | 114 | 7 | 133 | 4 | 137 | 6 | 0 | 0 | 69 | 5 | 2 | 0 |
| MaxQuant MBR+EBRCT | 119 | 12 | 124 | 13 | 130 | 20 | 1 | 1 | 91 | 2 | 28 | 1 |

**human-yeast**

| yeast spike-in protein concentration [%] | 10 vs 5 | | 10 vs 3.3 | | 5 vs 3.3 | |
|---|---|---|---|---|---|---|
| Method | tp | fp | tp | fp | tp | fp |
| **Quandenser+Triqler** | | | | | | |
| **- Tide** | **366** | **24** | **419** | **24** | **198** | **0** |
| - MODa | 223 | 11 | 296 | 15 | 77 | 0 |
| - Tide + MODa cascade | 334 | 13 | 382 | 20 | 154 | 0 |
| Tide+Triqler (without Quandenser/MBR) | 188 | 3 | 211 | 11 | 99 | 0 |
| MaxQuant MBR+Perseus | | | | | | |
| - $S_0 = 0.0$ | 226 | 36 | 266 | 32 | 133 | 22 |
| - $S_0 = 0.3$ | 322 | 34 | 361 | 27 | 72 | 5 |
| - $S_0 = 0.7$ | 289 | 27 | 358 | 25 | 1 | 0 |
| - $S_0 = 1.0$ | 188 | 18 | 356 | 21 | 1 | 0 |
| - original study | 296 | ? | 287 | ? | 107 | ? |
| MaxQuant MBR+EBRCT | 372 | 41 | 388 | 101 | 28 | 9 |

**BoxCar hela-ecoli**

| E. coli spike-in ratio [1 : $x$] | BoxCar 2 vs 12 | | | | Shotgun 2 vs 12 | | | |
|---|---|---|---|---|---|---|---|---|
| | 1-sided test | | 2-sided test | | 1-sided test | | 2-sided test | |
| **Method** | tp | fp | tp | fp | tp | fp | tp | fp |
| **Quandenser+Triqler** | | | | | | | | |
| **- Tide** | **977** | **7** | **977** | **32** | **964** | **10** | **964** | **218** |
| - MODa | 946 | 18 | 946 | 36 | 950 | 17 | 950 | 216 |
| - Tide + MODa cascade | 961 | 6 | 961 | 27 | 959 | 6 | 959 | 207 |
| Tide+Triqler (without Quandenser/MBR)* | 140 | 4 | 140 | 11 | 743 | 11 | 743 | 59 |
| MaxQuant MBR+Perseus** | | | | | | | | |
| - Student t-test $S_0 = 0.0$ | 965 | 16 | 971 | 3486 | 716 | 21 | 723 | 2228 |
| - Welch t-test $S_0 = 0.3$ | 972 | 19 | 975 | 2452 | 723 | 19 | 723 | 716 |

* BoxCar samples very few MS2 spectra; peptide identities are typically inferred with MBR to Shotgun runs, explaining the low sensitivity of the BoxCar runs without any form of MBR

** We used the MaxQuant output files from the original study where the data was searched against Swiss-Prot+TrEMBL, whereas we searched only against Swiss-Prot

Table S1: The table lists the number of true and false positive significantly differentially expressed proteins at a 5% reported FDR threshold for (1) Quandenser+Triqler, (2) for different values of $S_0$ for Welch's t-test in Perseus and (3) for the EBRCT method. Some results for Quandenser+Triqler and MaxQuant+EBRCT are left out here as they are listed in the main manuscript.

### 4 Posterior distributions for fold changes

We plotted the posterior distributions of fold changes as calculated by Triqler for the UPS-Yeast set (Figure S8) and the Human+Yeast and HeLa+Ecoli sets (Figure S9). We compared these to the results of MaxQuant+Perseus, where we instead of the posterior distribution plot the confidence interval using the mean and standard error of the mean.

Triqler achieves posterior estimates that approximate the true ratios quite well for a large proportion of the spiked-in proteins. Note that proteins with low FDR (towards the top of each panel) show fold changes very close to the expected value, but proteins with higher FDR (towards the bottom of each panel) show fold changes close to 0. The reason for the shift is that the proteins with higher FDR either have 1) peptides with high FDR, which have a harder time to overrule the prior, or 2) incorrect peptides that actually belong to the background. This is exactly the way that you would like a method to behave. You do not want to report inaccurately identified proteins as differentially abundant, i.e. having a fold change far from 0.

MaxQuant+Perseus achieves good quantitative accuracy and narrow confidence regions for highly confident proteins. However, we also see that many proteins for MaxQuant+Perseus have either large variance estimates, which often prevents $t$-tests from calling significance, or are not quantified because they had fewer than 2 peptides below 1% FDR.

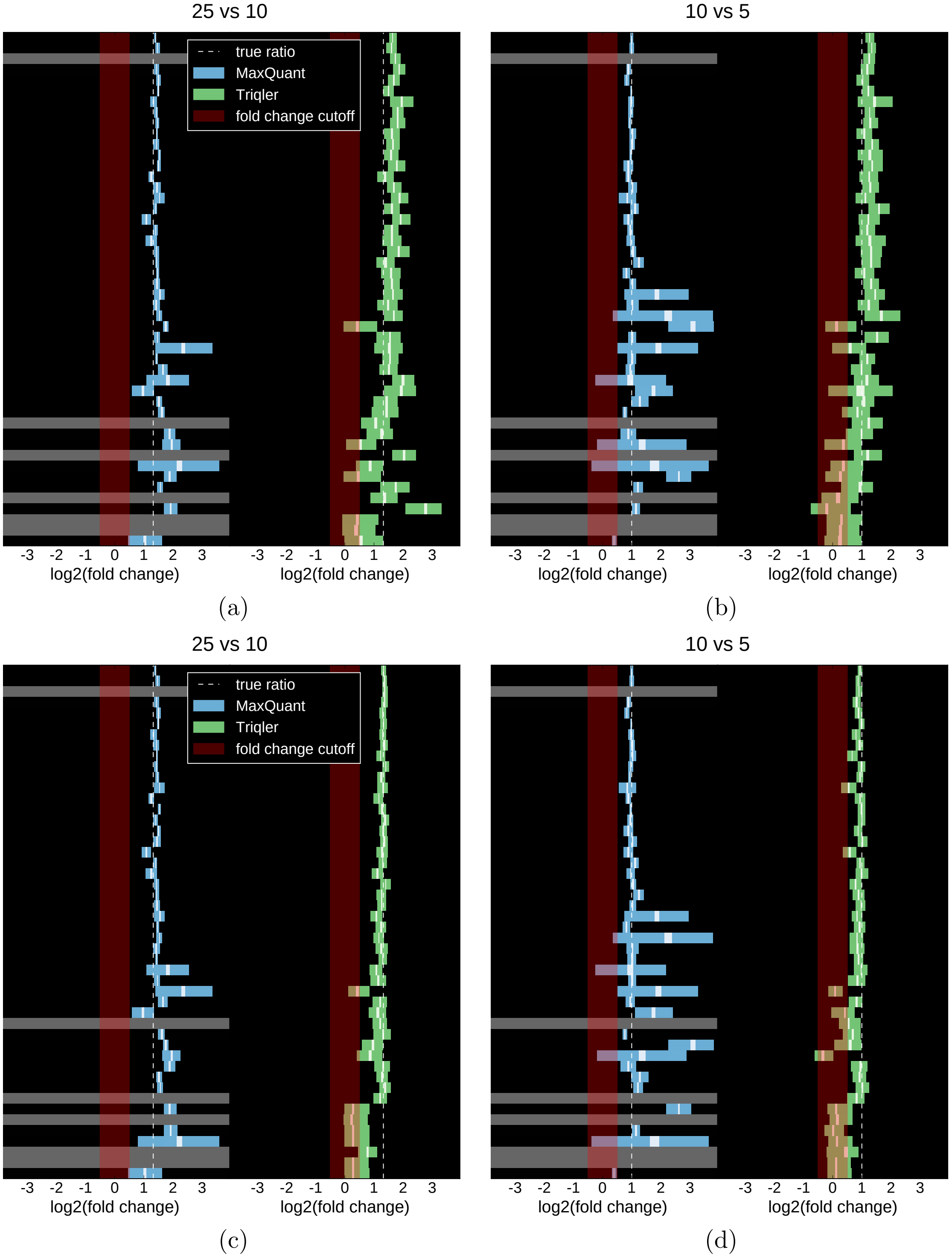


Figure S8: **The posterior distributions for the fold change of the spiked-in UPS proteins follow the true spike-in ratios well.** We compare the uncertainty of the fold change estimations of MaxQuant+Perseus (blue) and Quandenser+Triqler (green). Each row represents a UPS protein sorted from top to bottom by protein FDR, where the colored portions represent the 90% and the white portions the 10% confidence/credible intervals for the 25 vs 10 fmol **(a)** and 10 vs 5 fmol comparisons **(b)**. The quantitative accuracy of Quandenser+Triqler can be further improved by using the peak apex intensity, instead of the peak summed intensity, at the cost of reduced sensitivity **(c)**, **(d)**.

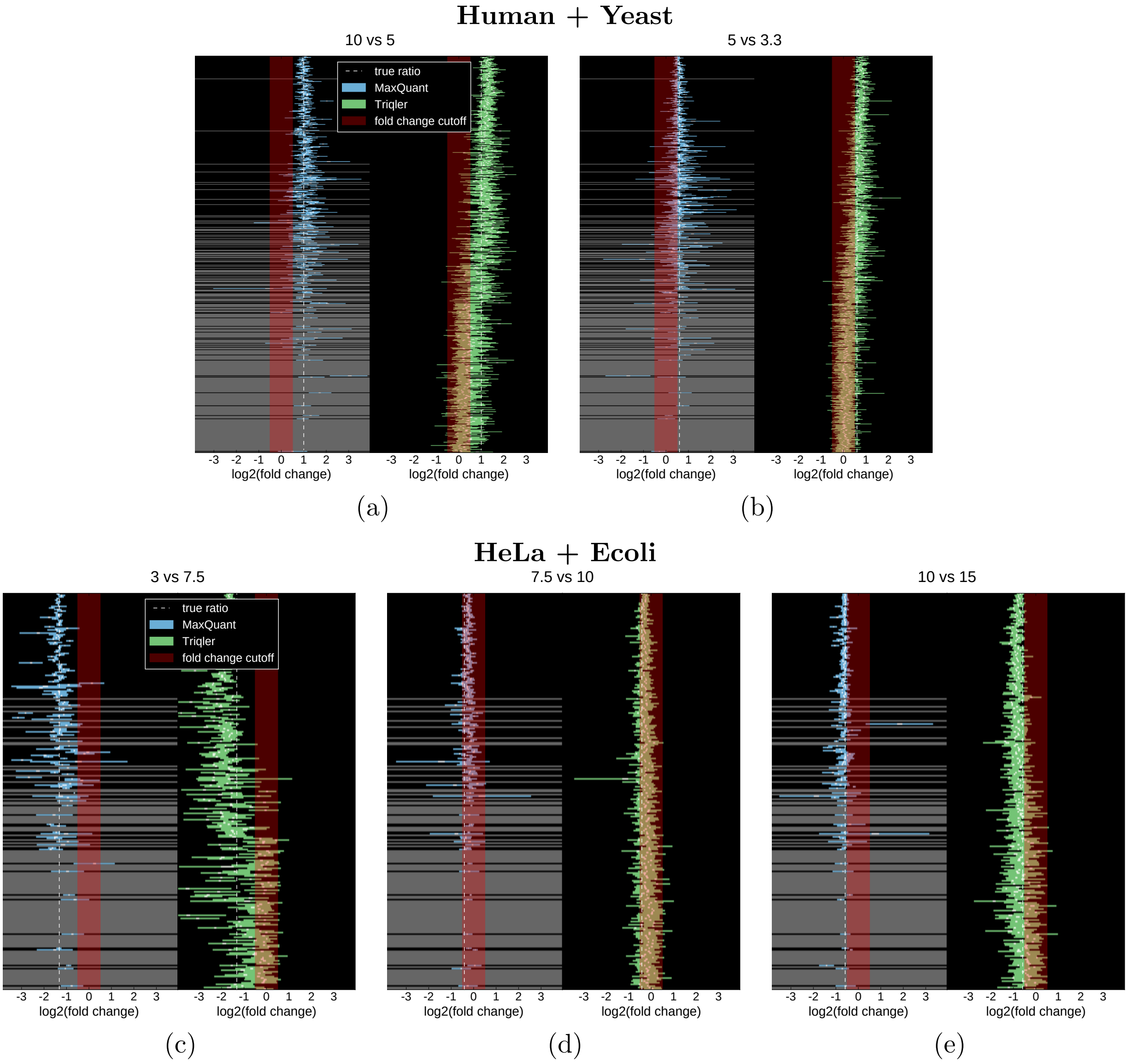


Figure S9: **Quandenser+Triqler provides stable estimates of the fold changes for both the human-yeast (a, b)and HeLa+Ecoli proteome mixture sets (c, d, e).** We compare the uncertainty of the fold change estimations of MaxQuant+Perseus (blue) and Quandenser+Triqler (green). Each row represents a, respectively, yeast or e. coli protein sorted from top to bottom by protein FDR, where the colored portions represent the 90% and the white portions the 10% confidence/credible intervals.

#### 5 Decoy features

We define an incorrect feature-feature match as a match where the two connected MS1 features do not originate from the same analyte. In a similar fashion to database searching, we model the incorrect matches by so-called decoy matches. Instead of reversing an amino acid sequence, we now ensure that the matched feature for a particular query feature is incorrect by searching in an m/z region offset by a fixed value of 5*1.000508 Th. The idea behind this offset is that the density, w.r.t. precursor m/z and retention time, of MS1 features is approximately the same in this offset region. This thus simulates the situation where a particular query feature is matched to a feature in the non-offset region if the truly matching feature is either not present, or if another incorrect feature is closer, by some weighting of the numeric features, such as precursor m/z and retention time difference.

To check the assumption that matches to decoy features mimic matches to incorrect features, we can plot the distribution of the numeric features and check if the distributions between target and decoy matches overlap in the region where we expect very few correct matches (Supplementary Figure S10). In our case, we see a good overlap between targets and decoys for large retention time and precursor m/z differences. For the precursor m/z differences, we observe a strange behavior in the $[(-)10, (-)50]$ ppm range, where there is a depletion of target features, which leads to conservative estimates of the FDR. We hypothesize this to be be due to peaks being aggregated within this window by the feature finding algorithm, Dinosaur.

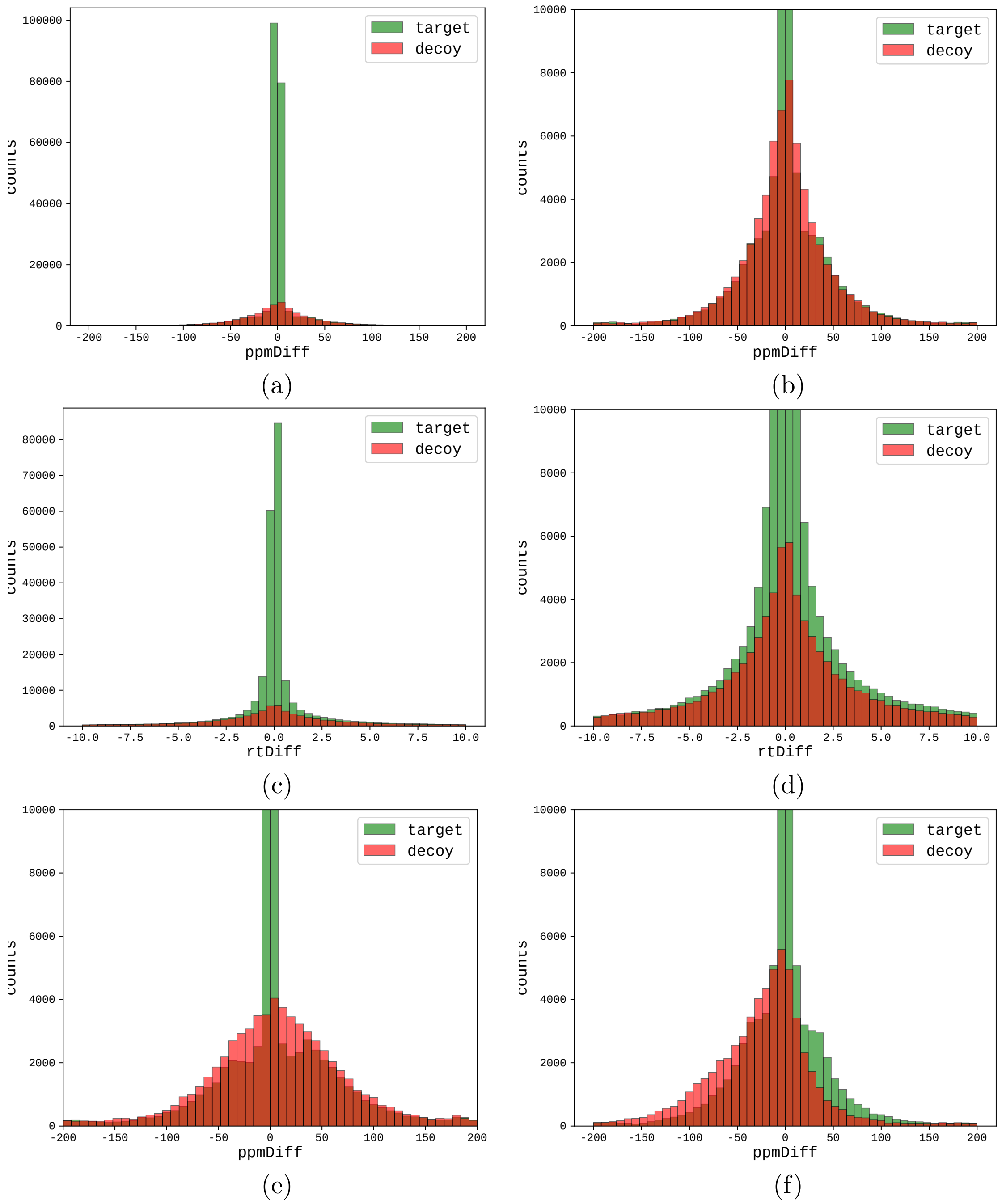


Figure S10: **Decoy MS1 features provide a good proxy for incorrect target MS1 features.** **(a)** Histogram of the precursor m/z difference in parts-per-million (ppm) for all matches in a single pair-wise alignment in the Hela-Ecoli mixture set. We see a clear enrichment of target MS1 features close to zero and good overlap between target and decoy feature matches far from zero. **(b)** same as (a), but zoomed in to 10000 counts. **(c)** same as (a), but for aligned retention time differences in minutes. Again, we see enrichment close to zero. **(d)** same as (c) but zoomed in to 10000 counts. Again, overlap of target and decoy feature matches for large retention time differences can be observed. **(e)** same as (b), but for Dinosaur in targeted mode. Again, we see good overlap from around 50 ppm. **(f)** same as (b), but using a decoy precursor m/z offset of 5.025 instead of $5 \cdot 1.000508$, mimicking the offset used in DeMix-Q. The non-integer shift of the precursor mass causes a bias in the mass difference distribution because peptide masses have to be additions of amino acids masses.

### 6 Functional annotation analysis

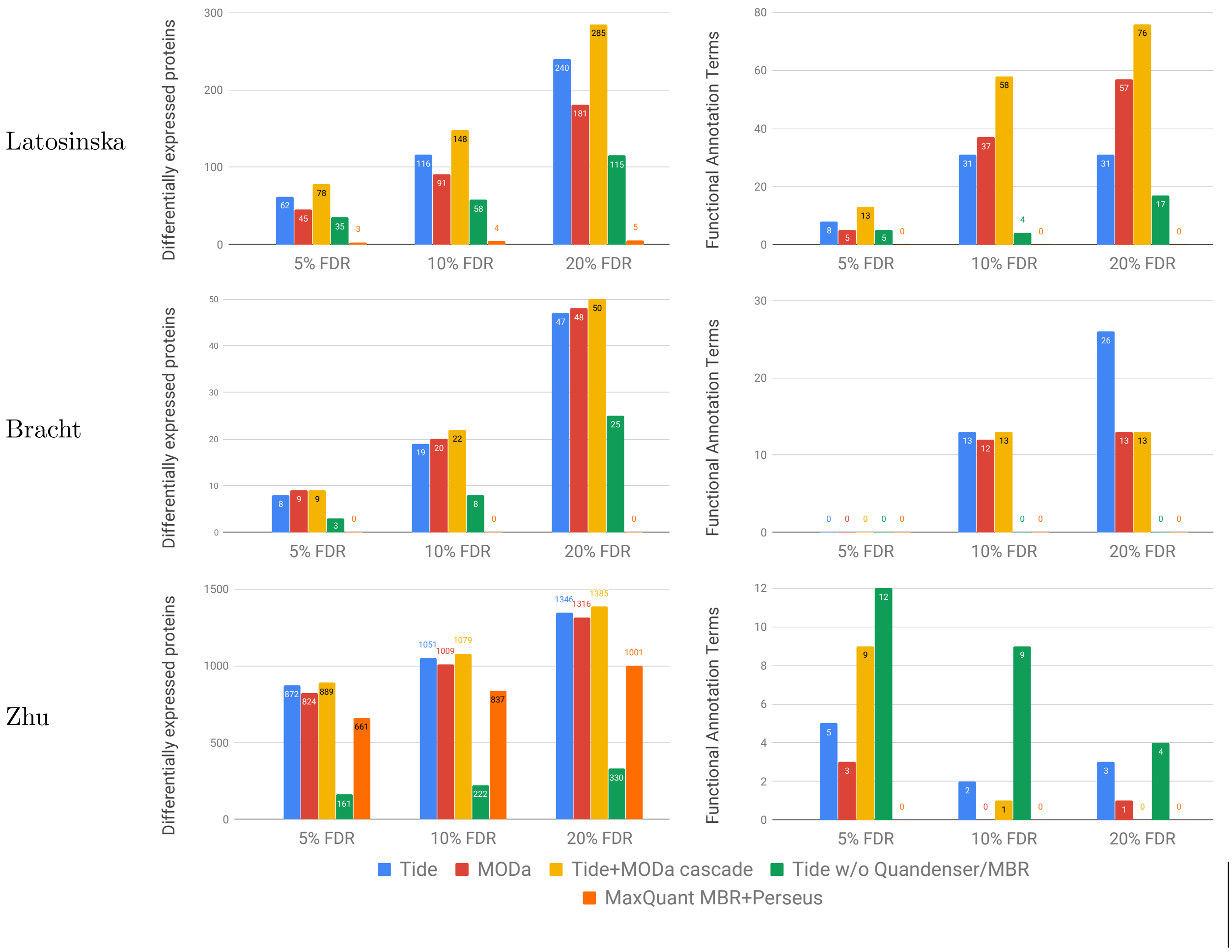


Figure S11: **Benchmark of differentially expressed proteins and enriched functional annotation terms.** The analyzed methods were three quantification-first approaches, Quandenser+Triqler when using the search engine Tide (**blue**), MODa (**red**), and a cascade search with first Tide and subsequently MODa (**yellow**); and three identification-first approaches, Tide and Triqler without clustering and the match-between-runs (MBR) feature (**green**), and MaxQuant with MBR followed by statistical analysis with Perseus (**orange**) and EBRCT (**light blue**). Overall, we discovered more differentially expressed proteins and enriched functional annotation terms with than without Quandenser. Notably, we found enriched functional annotation terms for the Bracht set for which no enrichments were previously found. The left plots show the number of differentially expressed proteins at 3 differential abundance FDR thresholds. The plots on the right show the number of significant functional annotation terms we discovered with DAVID using the sets obtained in the left plots. Note that the FDR reported in the plots on the right refer to the differential abundance FDR and not the functional annotation term FDR, which was kept fixed at 5%.

`Table_S2_largest_clusters_UPS`

Table S2: The table lists the largest unidentified spectrum clusters with their tentative peptide identifications in the UPS/Yeast dataser.

`Table_S3_FG_with_UPS_pattern`

Table S3: The table lists the feature groups with the smallest cosine distance to the UPS expression pattern with their tentative peptide identifications.

Table_S4_UPS_carbohydrate_frag

Table S4: The table lists the fragment ions of 5 consensus spectra that seem to come from carbohydrates instead of amino acid oligomers.

Table_S5_Latosinska_Tide_q5

Table S5: Significant enriched functional annotation terms for the Latosinska dataset at 5% differential expression FDR for Tide.

Table_S6_Latosinska_Tide_q10

Table S6: Significant enriched functional annotation terms for the Latosinska dataset at 10% differential expression FDR for Tide.

Table_S7_Latosinska_Tide_q20

Table S7: Significant enriched functional annotation terms for the Latosinska dataset at 20% differential expression FDR for Tide.

Table_S8_Latosinska_MODa_q5

Table S8: Significant enriched functional annotation terms for the Latosinska dataset at 5% differential expression FDR for MODa.

Table_S9_Latosinska_MODa_q10

Table S9: Significant enriched functional annotation terms for the Latosinska dataset at 10% differential expression FDR for MODa.

Table_S10_Latosinska_MODa_q20

Table S10: Significant enriched functional annotation terms for the Latosinska dataset at 20% differential expression FDR for MODa.

Table_S11_Latosinska_cascad_q5

Table S11: Significant enriched functional annotation terms for the Latosinska dataset at 5% differential expression FDR for a Tide followed by MODa cascade search.

Table_S12_Latosinska_cascad_q10

Table S12: Significant enriched functional annotation terms for the Latosinska dataset at 10% differential expression FDR for a Tide followed by MODa cascade search.

Table_S13_Latosinska_cascad_q20

Table S13: Significant enriched functional annotation terms for the Latosinska dataset at 20% differential expression FDR for a Tide followed by MODa cascade search.

Table_S14_Bracht_Tide_q5

Table S14: Significant enriched functional annotation terms for the Bracht dataset at 5% differential expression FDR for Tide.

Table_S15_Bracht_Tide_q10

Table S15: Significant enriched functional annotation terms for the Bracht dataset at 10% differential expression FDR for Tide.

Table_S16_Bracht_Tide_q20

Table S16: Significant enriched functional annotation terms for the Bracht dataset at 20% differential expression FDR for Tide.

Table_S17_Bracht_MODa_q5

Table S17: Significant enriched functional annotation terms for the Bracht dataset at 5% differential expression FDR for MODa.

Table_S18_Bracht_MODa_q10

Table S18: Significant enriched functional annotation terms for the Bracht dataset at 10% differential expression FDR for MODa.

Table_S19_Bracht_MODa_q20

Table S19: Significant enriched functional annotation terms for the Bracht dataset at 20% differential expression FDR for MODa.

Table_S20_Bracht_cascad_q5

Table S20: Significant enriched functional annotation terms for the Bracht dataset at 5% differential expression FDR for a Tide followed by MODa cascade search.

Table_S21_Bracht_cascad_q10

Table S21: Significant enriched functional annotation terms for the Bracht dataset at 10% differential expression FDR for a Tide followed by MODa cascade search.

Table_S22_Bracht_cascad_q20

Table S22: Significant enriched functional annotation terms for the Bracht dataset at 20% differential expression FDR for a Tide followed by MODa cascade search.

Table_S23_Zhu_Tide_q5

Table S23: Significant enriched functional annotation terms for the Zhu dataset at 5% differential expression FDR for Tide.

Table_S24_Zhu_Tide_q10

Table S24: Significant enriched functional annotation terms for the Zhu dataset at 10% differential expression FDR for Tide.

Table_S25_Zhu_Tide_q20

Table S25: Significant enriched functional annotation terms for the Zhu dataset at 20% differential expression FDR for Tide.

Table_S26_Zhu_MODa_q5

Table S26: Significant enriched functional annotation terms for the Zhu dataset at 5% differential expression FDR for MODa.

Table_S27_Zhu_MODa_q10

Table S27: Significant enriched functional annotation terms for the Zhu dataset at 10% differential expression FDR for MODa.

Table_S28_Zhu_MODa_q20

Table S28: Significant enriched functional annotation terms for the Zhu dataset at 20% differential expression FDR for MODa.

Table_S29_Zhu_cascad_q5

Table S29: Significant enriched functional annotation terms for the Zhu dataset at 5% differential expression FDR for a Tide followed by MODa cascade search.

Table_S30_Zhu_cascad_q10

Table S30: Significant enriched functional annotation terms for the Zhu dataset at 10% differential expression FDR for a Tide followed by MODa cascade search.

Table_S31_Zhu_cascad_q20

Table S31: Significant enriched functional annotation terms for the Zhu dataset at 20% differential expression FDR for a Tide followed by MODa cascade search.

### 7 Missing value analysis

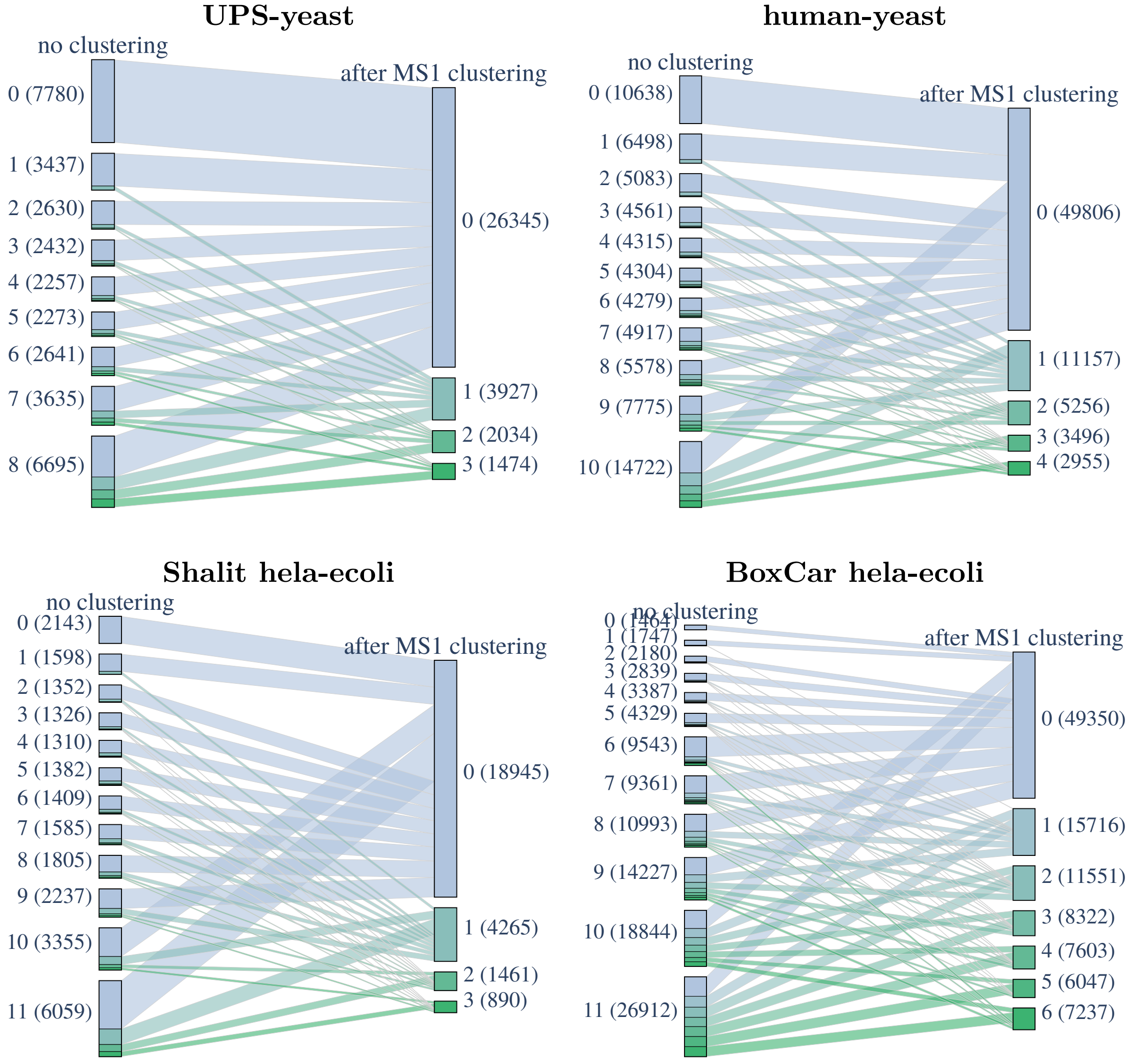


Figure S12: **MS1 clustering drastically reduces the number of missing values.** For each of the engineered datasets, we plot the number of feature groups with MS2 spectrum with $M$ missing values without clustering (left bars) and after MS1 clustering (right bars), annotated as "$M$ (number of feature groups)". We see that even feature groups with an MS2 spectrum in only a single run (the bottom bar on the left in each panel) can often be rescued by MS1 clustering, frequently even leading to no missing values at all.

**human-yeast**

| Method | 10 vs 5 | | 10 vs 3.3 | | 5 vs 3.3 | |
|---|---|---|---|---|---|---|
| | tp | fp | tp | fp | tp | fp |
| Quandenser+Tide+Triqler | | | | | | |
| - max-missing 8 | 434 | 43 | 511 | 54 | 296 | 2 |
| - max-missing 7 | 418 | 34 | 490 | 46 | 276 | 2 |
| - max-missing 6 | 401 | 32 | 465 | 40 | 251 | 1 |
| - max-missing 5 | 381 | 27 | 441 | 32 | 227 | 0 |
| **- max-missing 4** | 366 | 24 | 419 | 24 | 198 | 0 |
| - max-missing 3 | 343 | 19 | 395 | 14 | 177 | 0 |

Table S32: **Quandenser+Triqler achieves reasonable FDR control for the human-yeast set (11 runs in total), even with many allowed missing values.** The table lists the number of true and false positive significantly differentially expressed proteins at a 5% reported FDR threshold for different numbers of missing values for the human-yeast set using the Quandenser+Tide+Triqler pipeline. In the manuscript, max-missing 4 was used. As expected, sensitivity increases with more allowed missing values. The empirical FDR increases with higher number of allowed missing values as well and although it exceeds the reported 5% with more than 5 missing values, it stays below 10% even when allowing up to 8 missing values.

### 8 Analysis of increase in performance due to clustering

The inaccurately quantified peptides in Figure S13 could give rise to falsely quantified proteins, especially if their quantifications would be considered as reliable as peptides before MS1 matching. Of the feature groups that were added by MS1 matching 4124/18458 = 22% had a $|logFC| > 1.0$, for the feature groups before MS1 clustering, this rate was 2845/25681 = 11%. While the inaccurately quantified clusters might seem plentiful, by using our probabilistic graphical model, their influence is frequently down-weighted by considering matching probabilities and the agreement with other peptides of the protein. We indeed see a rise on protein level for the number of false positives (which are all human) from 3 to 14 after MS1 clustering for the 5vs10 comparison (Table S33), but the observed differential expression FDR stays below the reported 5%, rising from 1.6% to 3.9%.

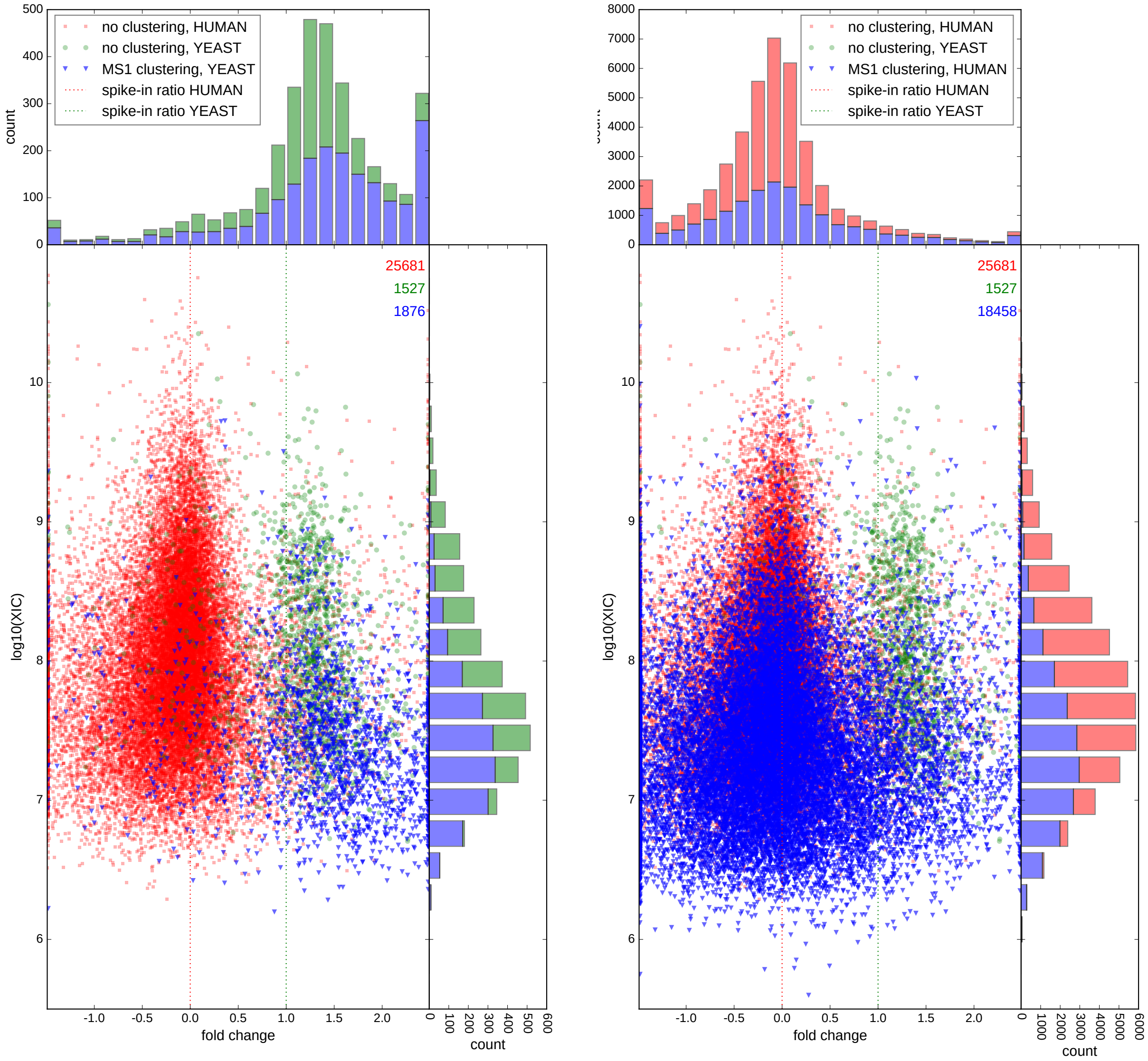


(a) (b)

Figure S13: **MS1 clustering (blue) almost doubles the number of peptides quantified at 1% FDR.** Scatter plots with histogram projections of peptide fold changes against extracted ion current (XIC) at 1% peptide-level identification FDR with at most 4 missing values for the human-yeast set. Fold changes are taken from predictions of the 10 vs 5 comparison, fold changes outside of the plotted range are set to the respective minimum or maximum fold change within the plotted range. We see a separation of human (red) and yeast peptides (green) before applying any clustering, which follow the spike-in ratios. We demonstrate that MS1 clustering more than doubles the number of yeast peptides **(a)** and adds around 70% more peptides for human **(b)**. These added peptides follow the expected fold changes and show a clear enrichment for lower XIC values, contributing to lowering the limit of detection (LOD).

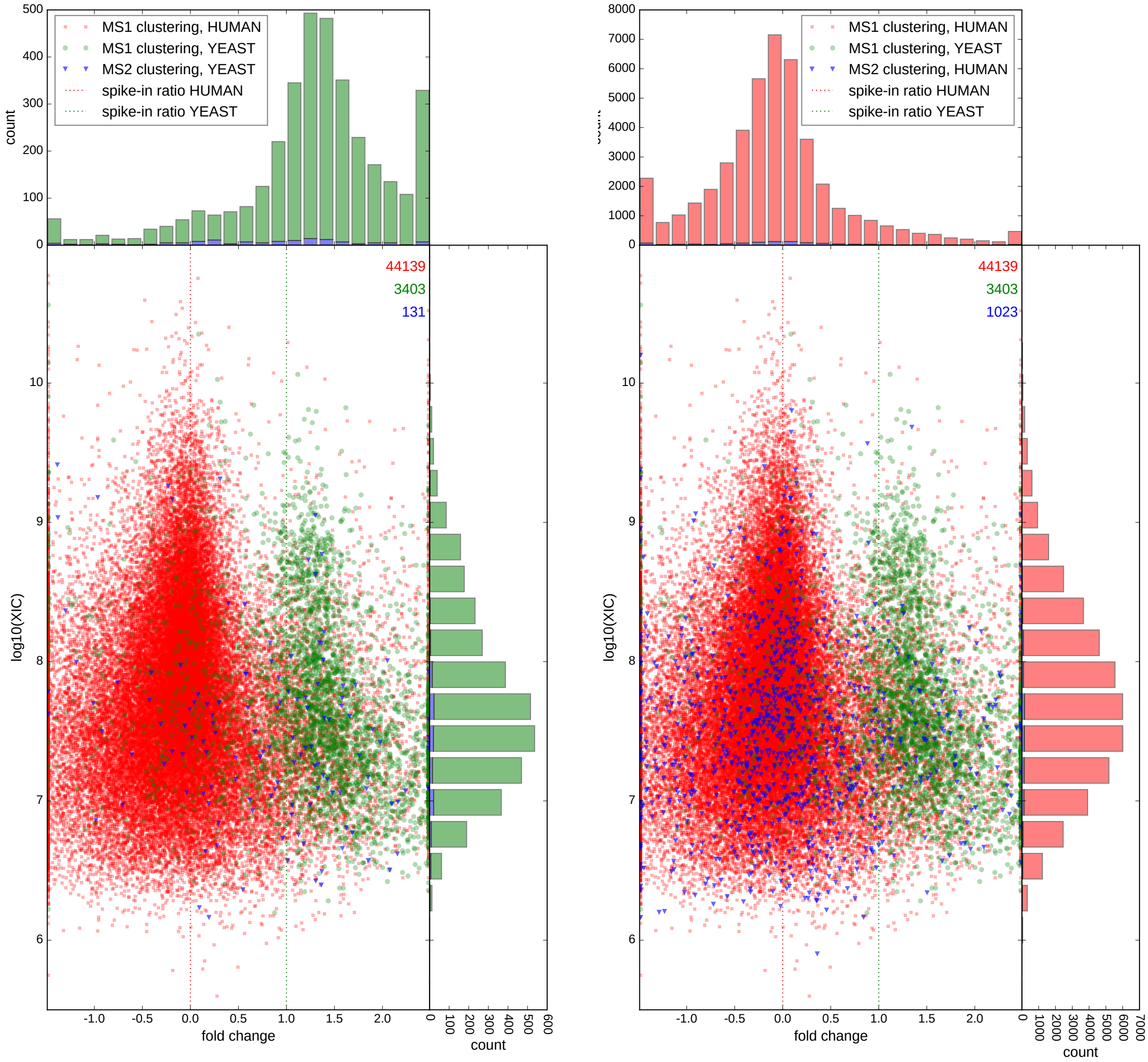


(a) (b)

Figure S14: **MS2 clustering (blue) only modestly increases the number of peptides quantified at** 1% **FDR.** Scatter plots with histogram projections of peptide fold changes against extracted ion current (XIC) at 1% peptide-level identification FDR with at most 4 missing values for the human-yeast set. Fold changes are taken from predictions of the 10 vs 5 comparison, fold changes outside of the plotted range are set to the respective minimum or maximum fold change within the plotted range. We see that human (red) and yeast peptides (green), with MS1 clustering but without MS2 clustering, follow the spike-in ratios. MS2 clustering only adds 4% more yeast peptides **(a)** and 2% more peptides for human **(b)**. Since MS1 clustering already allows identity propagation between features, the contribution of MS2 clustering is limited to reducing the number of hypotheses tested and increasing signal on noisy MS2 spectra through the formation of consensus spectra.

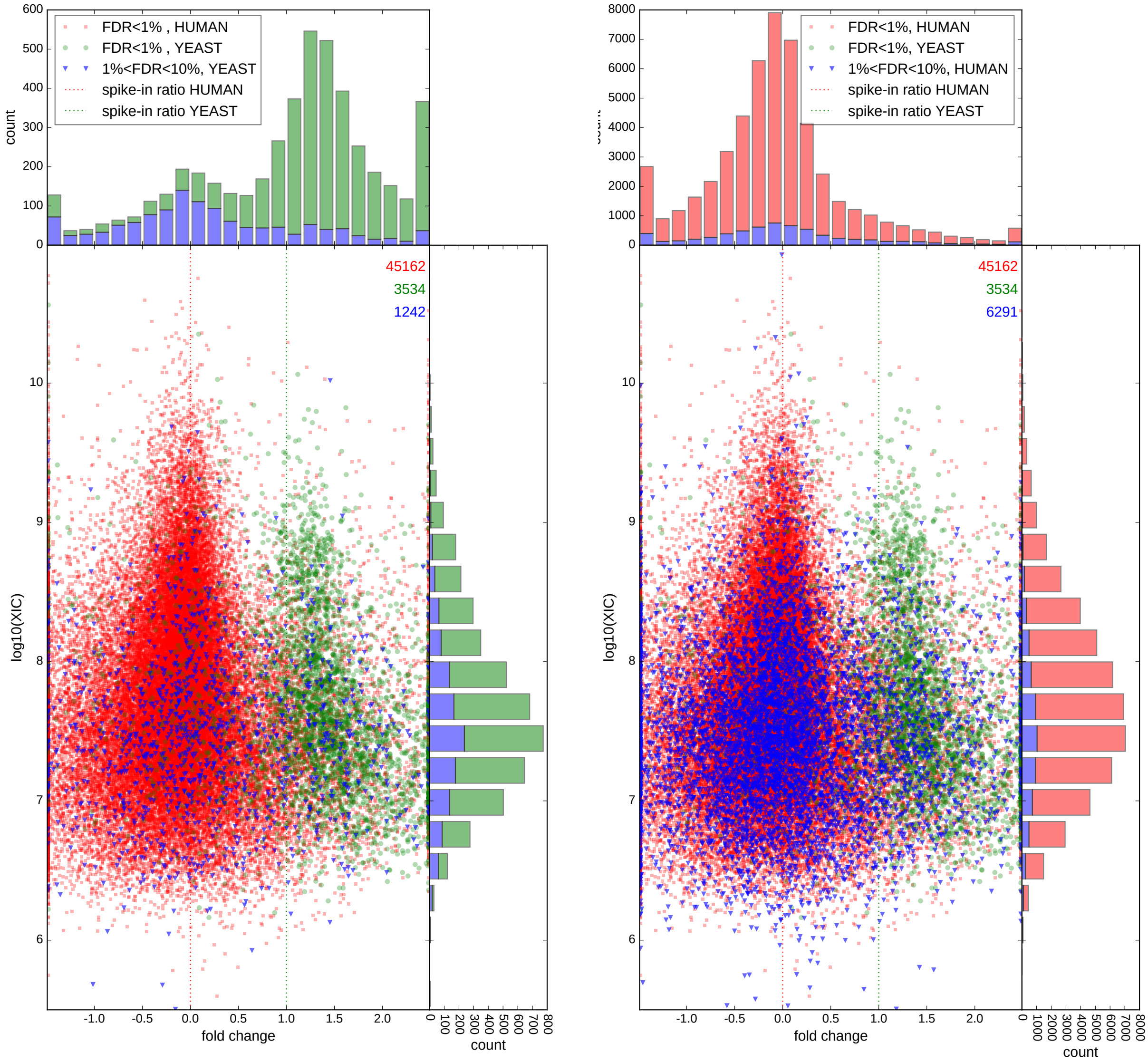


(a) (b)

Figure S15: **Peptide identification between 1% and 10% FDR (blue) could provide valuable extra information that is typically discarded.** Scatter plots with histogram projections of peptide fold changes against extracted ion current (XIC) at 1% peptide-level identification FDR with at most 4 missing values for the human-yeast set. Fold changes are taken from predictions of the 10 vs 5 comparison, fold changes outside of the plotted range are set to the respective minimum or maximum fold change within the plotted range. We see a separation of human (red) and yeast peptides (green), with MS1 and MS2 clustering, which follow the spike-in ratios. We see that although many peptides identified as coming from yeast between 1% and 10% FDR are most likely misidentified as they follow the human spike-in ratio, a non-negligible amount of peptides do follow the correct spike-in ratio of yeast as well **(a)**. For the majority class of human, this looks better on first sight, but will probably still contain many false positives **(b)**. Even though the peptide identification between 1% and 10% FDR should be handled with the greatest caution, they could help to confirm or oppose evidence below the 1% FDR threshold.

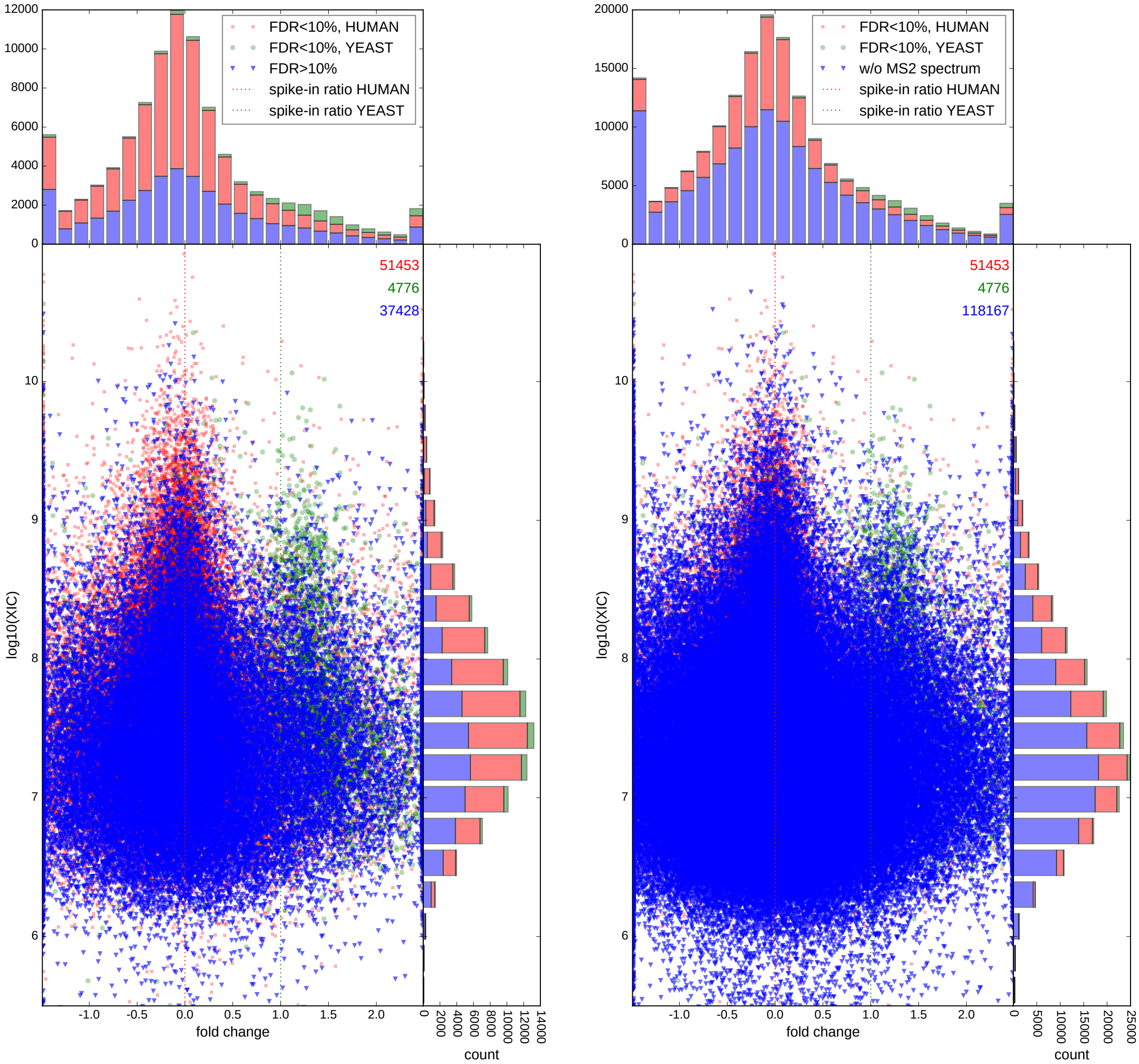


(a) (b)

Figure S16: **MS1 feature groups without identification (left, blue) and without MS2 spectrum (right, blue) represent a sizeable part of the proteome that remains unknown.** Scatter plots with histogram projections of peptide fold changes against extracted ion current (XIC) at 10% peptide-level identification FDR with at most 4 missing values for the human-yeast set. Fold changes are taken from predictions of the 10 vs 5 comparison, fold changes outside of the plotted range are set to the respective minimum or maximum fold change within the plotted range. We see a separation of human (red) and yeast peptides (green), with MS1 and MS2 clustering, which follow the spike-in ratios. MS1 feature groups without identification at 10% FDR still make up around 40% of the data **(a)**. MS1 feature groups without MS2 spectrum almost triples the data, which in the future might be identifiable by, e.g. accurate mass and retention time predictions **(b)**.

**UPS-yeast**

| Method | Num feature groups | | 25 vs 10 | | 25 vs 5 | | 10 vs 5 | |
|---|---|---|---|---|---|---|---|---|
| | UPS | yeast | tp | fp | tp | fp | tp | fp |
| No clustering, Tide, FDR < 1% | 230 | 9193 | 37 | 0 | 38 | 0 | 33 | 1 |
| + MS1 clustering | (+139) 369 | (+2956) 12149 | 44 | 2 | 46 | 2 | 39 | 0 |
| + MS2 clustering | (+4) 373 | (+322) 12471 | 43 | 2 | 45 | 2 | 38 | 0 |
| + 1% < FDR < 10% | (+50) 423 | (+3115) 15595 | 43 | 2 | 46 | 2 | 38 | 2 |
| + FDR > 10% | (+34166) 50184 | | 43 | 1 | 46 | 0 | 37 | 0 |
| + no MS2 spectrum | (+70542) 120726 | | - | - | - | - | - | - |

**human-yeast**

| Method | Num feature groups | | 10 vs 5 | | 10 vs 3.3 | | 5 vs 3.3 | |
|---|---|---|---|---|---|---|---|---|
| | yeast | human | tp | fp | tp | fp | tp | fp |
| No clustering, Tide, FDR < 1% | 1527 | 25681 | 182 | 3 | 205 | 1 | 85 | 0 |
| + MS1 clustering | (+1876) 3403 | (+18458) 44139 | 346 | 14 | 385 | 21 | 176 | 0 |
| + MS2 clustering | (+131) 3534 | (+1023) 45162 | 346 | 16 | 388 | 22 | 178 | 0 |
| + 1% < FDR < 10% | (+1242) 4776 | (+6291) 51453 | 366 | 24 | 419 | 24 | 198 | 0 |
| + FDR > 10% | (+37428) 93657 | | 366 | 24 | 419 | 24 | 198 | 0 |
| + no MS2 spectrum | (+118167) 211824 | | - | - | - | - | - | - |

**Shalit hela-ecoli**

| Method | Num feature groups | | 3 vs 7.5 | | 3 vs 10 | | 3 vs 15 | | 7.5 vs 10 | | 7.5 vs 15 | | 10 vs 15 | |
|---|---|---|---|---|---|---|---|---|---|---|---|---|---|---|
| | E. coli | HeLa | tp | fp | tp | fp | tp | fp | tp | fp | tp | fp | tp | fp |
| No clustering, Tide, FDR < 1% | 269 | 5836 | 44 | 0 | 47 | 0 | 52 | 0 | 0 | 0 | 48 | 0 | 41 | 0 |
| + MS1 clustering | (+796) 1065 | (+7625) 13461 | 138 | 0 | 146 | 1 | 176 | 0 | 0 | 0 | 161 | 0 | 112 | 0 |
| + MS2 clustering | (+35) 1100 | (+613) 14074 | 142 | 0 | 150 | 1 | 183 | 0 | 0 | 0 | 167 | 0 | 116 | 0 |
| + 1% < FDR < 10% | (+399) 1499 | (+4551) 18625 | 159 | 2 | 167 | 2 | 206 | 0 | 0 | 0 | 185 | 1 | 137 | 0 |
| + FDR > 10% | (+38232) 58356 | | 190 | 7 | 198 | 4 | 229 | 0 | 0 | 0 | 194 | 3 | 138 | 0 |
| + no MS2 spectrum | (+156684) 215040 | | - | - | - | - | - | - | - | - | - | - | - | - |

**BoxCar hela-ecoli***

| Method | Num feature groups | | BoxCar 2 vs 12 | | | | Shotgun 2 vs 12 | | | |
|---|---|---|---|---|---|---|---|---|---|---|
| | | | 1-sided test | | 2-sided test | | 1-sided test | | 2-sided test | |
| | E. coli | HeLa | tp | fp | tp | fp | tp | fp | tp | fp |
| No clustering, Tide, FDR < 1% | 2626 | 17269 | 460 | 14 | 460 | 21 | 475 | 8 | 475 | 40 |
| + MS1 clustering | (+6665) 9291 | (+34324) 51593 | 999 | 14 | 999 | 54 | 977 | 3 | 977 | 176 |
| + MS2 clustering | (+262) 9553 | (+1543) 53136 | 1007 | 14 | 1007 | 53 | 982 | 3 | 982 | 186 |
| + 1% < FDR < 10% | (+1486) 11039 | (+10735) 63871 | 1027 | 18 | 1027 | 68 | 999 | 3 | 999 | 216 |
| + FDR > 10% | (+62175) 137085 | | 1029 | 18 | 1029 | 71 | 1002 | 4 | 1002 | 234 |
| + no MS2 spectrum | (+229680) 366765 | | - | - | - | - | - | - | - | - |

* As the BoxCar runs rely on identification propagation from the Shotgun runs, no representative results could be generated from the separate analyses, as was done in the rest of the manuscript. Therefore, results are shown for a combined analysis of the 6 BoxCar with the 6 Shotgun runs with maximum allowed missing values, $M = 6$.

Table S33: **MS1 clustering is the biggest contributor to the increased sensitivity of our pipeline.** By filtering the feature groups that are added by certain steps of our pipeline, we could look at the contributions of these steps. MS1 clustering clearly shows the biggest contribution to sensitivity, whereas contributions of MS2 clustering and including less confident peptides is rather modest.

#### 9 Summary of datasets

$M$: max missing values
$fc_{eval}$: $|\log_2|$ fold change eval for Triqler
MS2 sp: number of MS2 spectra
SFP: number of spectrum-feature pairs, number of spectra that need to be searched by the search engine
FG: feature groups
FG sp: number of feature groups with at least one MS2 consensus spectrum
cs sp.: number of consensus spectra
cs SFP: consensus spectrum-feature pairs, number of spectra that need to be searched
rd SFP: reduction in number of spectrum-feature pairs that need to be searched

| Dataset | PRIDE ID | runs | $M$ | $fc_{eval}$ | MS2 sp | SFP | FG | FG sp | cs sp | cs SFP | rd SFP |
|---|---|---|---|---|---|---|---|---|---|---|---|
| UPS-yeast | PXD002370 | 9 | 3 | 0.8 | 535k | 934k | 121k | 51k | 61k | 109k | 88% |
| human-yeast | PXD007683 | 11 | 4 | 0.5 | 807k | 1.12M | 212k | 94k | 116k | 178k | 84% |
| Shalit hela-ecoli | PXD001385 | 12 | 3 | 0.5 | 340k | 533k | 215k | 53k | 67k | 96k | 82% |
| BoxCar hela-ecoli | PXD006109 | 12 | 6 | 0.5 | 629k | 1.02M | 370k | 124k | 134k | 218k | 79% |
| Latosinska | PXD002170 | 8 | 4 | 0.8 | 413k | 991k | 83k | 47k | 122k | 183k | 82% |
| Bracht | PXD001474 | 27 | 7 | 0.5 | 1.01M | 1.47M | 69k | 45k | 106k | 150k | 90% |
| Zhu | PXD006847 | 18 | 11 | 1.0 | 593k | 1.02M | 73k | 52k | 117k | 187k | 82% |

Table S34: **Summary of the datasets, including used parameters and reduction in searched MS2 spectra.**
